# Oxonium Ion-Guided Ion Mobility-Assisted Glycoproteomics on the timsTOF Pro

**DOI:** 10.1101/2022.07.04.498688

**Authors:** Soumya Mukherjee, Andris Jankevics, Florian Busch, Markus Lubeck, Yang Zou, Gary Kruppa, Albert J. R. Heck, Richard A. Scheltema, Karli R. Reiding

## Abstract

Spatial separation of ions in the gas-phase, providing information about their size as collisional cross-sections, can readily be achieved through ion mobility. The timsTOF Pro series combines a trapped ion mobility device with a quadrupole, collision-cell and a time-of-flight analyser to enable the analysis of ions at great speed. Here, we show that the timsTOF Pro is capable of physically separating *N*-glycopeptides from non-modified peptides and producing high-quality fragmentation spectra, both beneficial for glycoproteomics analyses of complex samples. The glycan moieties enlarge the size of glycopeptides compared to non-modified peptides, yielding a clear cluster in the mobilogram that, next to increased dynamic range from the physical separation of glycopeptides and non-modified peptides, can be used to make an effective selection filter for directing the mass spectrometer to analytes of interest. This new approach was applied to selected glycoproteins, human plasma- and neutrophil-derived glycopeptides. We show that the achieved physical separation, combined with the focussing of the mass spectrometer, allows for improved extraction of information from the samples, even at shorter LC gradients of 15 min. We validated our approach on human neutrophil and plasma samples of known make-up, in which we captured the anticipated glycan heterogeneity (paucimannose, phosphomannose, high mannose, hybrid and complex glycans) from plasma and neutrophil samples at the expected abundances. As the method is compatible with off-the-shelve data acquisition routines and data analysis software, it can readily be applied by any laboratory with a timsTOF Pro and is reproducible as demonstrated by a comparison between two laboratories.

---

Protein glycosylation is a highly abundant co- and post-translational modification (PTM), in which glycan moieties of varying complexity are covalently attached to specific residues in proteins(1). Protein glycosylation plays diverse roles in biological systems, influencing processes such as cell-cell adhesion, immunity and signalling through cellular recognition(2). Glycans most frequently attach to the proteins either via *N*-glycosidic linkages to asparagine residues (*N*-glycans) or via *O*-glycosidic linkages to the serine or threonine residues (*O*-glycans)(3, 4). A single glycoprotein is known to exhibit multiple glycoforms, displayed by both glycan micro-heterogeneity (different oligosaccharides can attach at the same site) and macro-heterogeneity (glycosylation site occupancy) per site, while sites across a given protein can be differentially regulated as well, i.e., meta-heterogeneity(5). Alterations in these glycosylation patterns have been well-documented between physiological and disease states(6, 7). Due to its biological importance, being dynamically regulated in response to any changes in homeostasis, glycosylation is an important target in biomarker research and biopharmaceutical development(8–11). This emphasizes the need of highly sensitive and precise analytical tools that can identify the highly diverse glycosylation patterns and localize them site-specifically on the proteins they adorn.

Mass spectrometric detection of glycans and intact glycopeptides has emerged as an attractive glycoproteomics analytical platform. Recent progress in workflows, including glycopeptide extraction/enrichment, hybrid mass spectrometric fragmentation, and data analysis, have made detection of glycopeptides increasingly achievable(12–17). Notwithstanding these advances over the past decade, characterization and quantitation of intact glycopeptides from complex datasets remains a bottleneck due to their inherent glycan heterogeneity, ionization and separation characteristics, and their relative low abundance compared to non-modified peptide counterparts(18). Optimised methods are clearly needed.

Ion mobility (IM) devices can separate ions by their collisional cross-section (CCS, Ω) at high speed (typically in the order of 10-100 ms)(19–21). Such devices typically are employed between liquid chromatography (LC) and the mass analyzer to provide an extra level of separation for the molecules of interest and provide improved dynamic range for the mass analysis. For this to work efficiently, a high-speed mass analyser is required, making TOF analysers attractive as they can operate at a scan-rate in the range of 100 kHz and thus can efficiently sample the ions eluting from the IM. Of the different conceptual devices to achieve gas-phase fractionation, trapped ion mobility separation (TIMS) can be packaged in a small device only requiring low operating voltages and providing efficient ion usage. In this device, ions are balanced in a constant gas stream by an electrical field allowing them to be stored at different positions. The ions can then be eluted – ordered by low mobility with large CCS to high mobility with small CCS – by lowering the electrical potential after which they are subsequently transferred to the mass analyzer. The timsTOF Pro (Bruker Daltonics) makes this effective combination and was recently shown to provide high analyte coverages in proteomic, lipidomic and metabolomic studies(22–25). With the data acquisition approach parallel accumulation serial fragmentation (PASEF) this instrument is capable to separate and accurately detect biological molecules (peptides, lipids, metabolites) at very fast scan rates(26, 27).

Recently, ion mobility separation mass spectrometry (IM-MS) emerged as a promising tool for characterizing glycosylated species (28–31). Glycopeptides, due to their inherent physical properties, have been shown to typically separate from non-modified peptides within both drift-tube and traveling-wave ion mobility mass spectrometers(32, 33). This enabled glycopeptides to be isolated in individual mobility windows with lower peptide components and chemical noise. This increases the signal/noise ratio, which is essential for improved detection of low abundant glycopeptide ions. Encouraged by this previous work, we hypothesized that, due to their inherent physical properties, *N*-glycopeptides have different mobility and would therefore cluster in a specific ion mobility region inside the TIMS device that is distinct from non-modified peptides (Figure 1). In this work, we first optimize the fragmentation settings with two purified glycoproteins for high quality fragmentation spectra boasting highly visible diagnostic glycan fragments, i.e., the glycan oxonium ions, as well as highly informative peptide backbone cleavages that can be used to confidently identify both the peptides and the attached glycan moieties. In addition, we demonstrate that the *N*-glycopeptides indeed cluster in a specific ion mobility region that is distinct from the localisation of non-modified peptides, and that physical separation of the two classes of molecules can be achieved. We validated the region of interest (ROI) of glycopeptides in the timsTOF Pro using two biological sample of higher complexity, enzymatic digests of human neutrophils and human plasma, to characterize the ion mobility space occupied by the heterogenous *N*-glycopeptides. The combination of these optimisations on the PASEF method led to a glycoproteomics method capable of identifying diverse and heterogeneous *N*-glycopeptides at both high-confidence and high-throughput on the timsTOF Pro.

**Figure 1.**
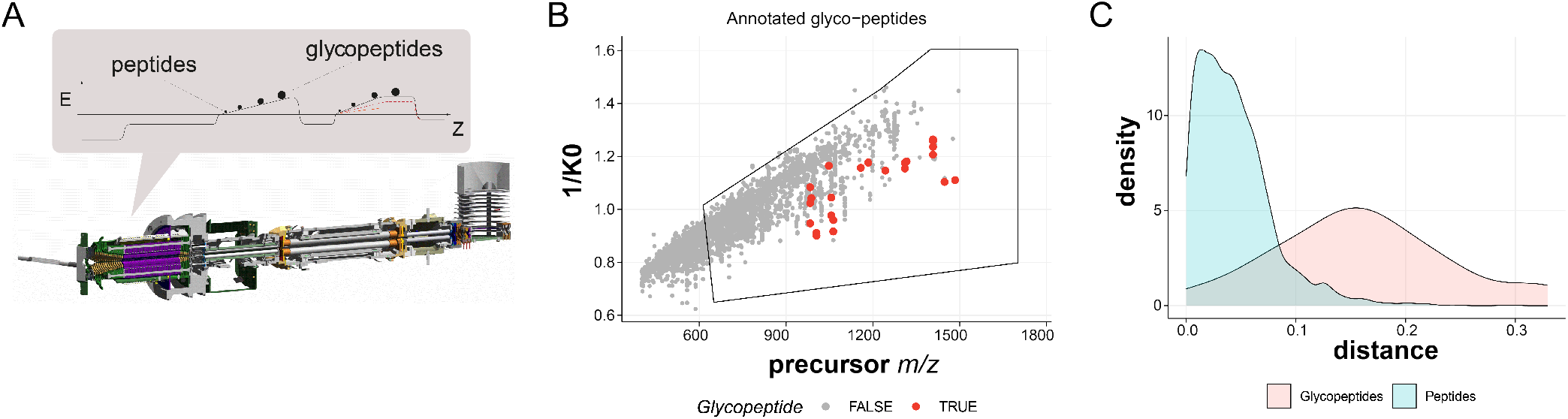
Ion Mobility-Assisted Glycoproteomics on the timsTOF Pro. **(A)** Schematic representation of the instrument with the conceptual operation of the TIMS device separation of the glycopeptides from the non-modified peptides. **(B)** Distribution of the ion signals in *m/z vs*. ion mobility (1/K0) for all classes of ions including non-modified peptides (gray) vs. glycopeptides (red). A schematic polygon is shown that encompasses the glycopeptide ion cluster in the ion mobility domain. (C) Density diagram displaying the physical separation of the glycopeptides from unmodified peptides in the ion mobility space (squared distance from linear fit of all data points in the data set).

## Materials and methods

### Chemicals and materials

Sodium deoxycholate (SDC), tris (2-carboxyethyl)phosphine (TCEP), Tris (tris (hydroxymethyl) aminomethane), CAA (chloroacetamide), NaOH (sodium hydroxide), TFA (trifluoroacetic acid) were purchased from Sigma-Aldrich (Steinheim, Germany). Formic acid (FA) was purchased from Merck (Darmstadt, Germany). Acetonitrile (ACN) was purchased from Biosolve (Valkenswaard, The Netherlands). Oasis µElution HLB and PRiME HLB plates were purchased from Waters (Etten-Leur, the Netherlands). Milli-Q was produced by an in-house system (Millipore, Billerica, MA). Both phosphoSTOP and cOmplete Mini EDTA-free were purchased from Roche (Woerden, the Netherlands). GluC was obtained from Roche (Indianapolis, IN). Recombinant tissue non-specific alkaline phosphatase (TNAP) was a gift from Copenhagen Centre of Glycomics. Histidine rich glycoprotein (HRG) was purified from human plasma with a cobalt-loaded resin (Thermo Scientific, Waltham, America) using IMAC-based enrichment (56). Commercial sialylglycopeptide (α2,6-SGP) and asialo-SGP were purchased from Fushimi Pharmaceutical Co., Ltd. Commercial pooled human plasma was purchased from Affinity Biologicals (Ancaster, ON). Purified human neutrophils, prepared as described previously(34), were a kind gift from the Department of Molecular and Cellular Homeostasis, Sanquin Research, Amsterdam, The Netherlands.

### Proteolytic digestion of nondepleted human plasma

10 µL of pooled human plasma was mixed with 50 volumes of SDC buffer (1% SDC, 50 mM Tris-HCl (pH 8.5), 10 mM TCEP, 30 mM CAA and boiled for 10 minutes at 95 °C. The samples were cooled and digested with a combination of Lys-C (1:75 enzyme to protein) for 4 hours followed by trypsin (1:20 enzyme to human plasma and 1:35 enzyme to HRG) at 37 °C overnight. The samples were quenched with 10% TFA to a final concentration of 1% TFA (0.5% TFA for HRG) and SDC was precipitated after centrifugation at 14,000 rpm for 10 min. The supernatant was transferred to a new tube and desalted using the µElution HLB plate. The desalted samples were lyophilized and stored at -80 °C before MS analysis.

### Cell lysis and proteolytic digestion of human neutrophils

The pooled human neutrophils from healthy donors were enriched by Percoll (GE Healthcare, Sweden) density gradient centrifugation as previously described(34). Neutrophil cell pellets were resuspended in 100 µL lysis buffer containing 100 mM Tris-HCl (pH 8.5), 7 M urea, 5 mM TCEP, 30 mM CAA, Triton X-100 (1%), 2 mM magnesium sulfate, phosphoSTOP and cOmplete Mini EDTA-free protease inhibitor. Then, cells were disrupted by sonication for 10 min (alternating 20 s off and 40 s off) using a Bioruptor Plus (Diagenode, Seraing, Belgium). Cell debris was removed by centrifugation at 14,000 rpm for 1 h at 4 °C and the supernatant was retained. Impurities were removed by methanol/chloroform protein precipitation as follows: 100 µL of supernatant was mixed with 400 µL of methanol, 100 µL chloroform and 300 µL of ultrapure water with thorough vortexing after each addition. The mixture was then centrifuged for 10 min at 5000 rpm at room temperature. The upper layer was discarded, and 300 µL of methanol was added. After sonication and centrifugation (5000 rpm, 10 min at room temperature), the solvent was removed, and the precipitate was allowed to dry in air inside a chemical hood. The pellet was resuspended in SDC buffer. GluC was then added to digest proteins for 3 h at an enzyme to protein ratio of 1:75 w/w at 37 °C. The resulting peptide mixtures were further digested overnight at 37 °C trypsin (1:20 w/w enzyme to protein ratio). The next day the SDC was precipitated via acidification to a final 0.5% TFA concentration. The peptides in the supernatant were desalted using an Oasis PRiME HLB plate and lyophilized and stored at – 80 °C prior to MS analysis.

### Data Acquisition

Tryptic peptides originating from the individual purified glycoproteins, as well as the more complex biological samples, were separated by using an Ultimate 3000 nanoUHPLC (Thermo Fischer Scientific) coupled on-line to a timsTOF Pro mass spectrometer (Bruker Daltonik). Peptides and glycopeptides were analytically separated on an Ion Optics nanoUHPLC column (75 µm × 25 cm, 1.6 µm, C18; Ion Optics, Australia), heated to 50 °C at a flow rate of 400 nL/min. LC mobile phases A and B were water with 0.1% formic acid (v/v) and ACN with formic acid 0.1% (v/v), respectively. The nanoLC was coupled to the timsTOF Pro via a modified nanoelectrospray ion source (Captive Spray, Bruker Daltonik). Initially, we used a 120 min gradient for the purified glycoprotein samples, while the plasma and neutrophil glycoprotein samples were separated using a 150 min gradient. The SGP and asialo-SGP were separated using a 15 min gradient. In the 120 min experiments the gradient was kept at 1% B for 13 min, increased to 3% B over the next 0.1 min, followed by an increase from 3% to 30% B over 90 min. For column wash, solvent B concentration was increased to 30% to 35% within 5 min and then at 80% for a further 1 min and kept at that concentration for an additional 5 min followed by re-equilibration to buffer A for 5 min. In the 180 min (150 min gradient) experiments used for the complex glycoprotein samples, and similarly for experiments at different gradient lengths at 60 min, 30 min and 15 min, the time between 3 and 30% B was modified accordingly.

Data acquisition on the timsTOF Pro was performed using otof-Control 6.0. Starting from the PASEF method optimized for standard proteomics(26), we integrated the glycan-specific polygon (as depicted in the figures). The following parameters were adapted. For the CaptiveSpray source inlet, the capillary voltage was set to 1500 V. The nebulizer dry gas flow rate was set to 3 L/min at 180 °C. TIMS region voltages were optimized at -20, -160, 110, 110, 0, and 75 V for Δ1-Δ6, respectively. TIMS RF was set to 350 Vpp. The allowed charge states for PASEF precursors were restricted to 2-5. The precursor intensity threshold was set to a target value of 20,000 counts, with dynamic exclusion release after 0.4 min. Each PASEF MS/MS frame consisted of two merged TIMS scans acquired for low and high collision energy profile for glycan specific ions in the stepped collision energy (PASEF SCE) method. In the first frame the base values for mobility dependent collision energy ramping were set to 100 eV at an inverse reduced mobility (1/K0) of 1.60 V.s/cm^2^ and 40 eV at 0.6 V.s/cm^2^, while in the second frame mobility dependent collision energy ramping were set to 65 eV at an inverse reduced mobility (1/K0) of 1.60 V.s/cm^2^ and 35 eV at 0.6 V.s/cm^2^. Collision energies were linearly interpolated between these two 1/K0 values and kept constant above or below these base points (see “Results and Discussion” for more details). PASEF without stepping consisted of only one TIMS scan with mobility dependent collision energy ramping set at 59 eV from reduced mobility (1/K0) of 1.60 V.s/cm^2^ to 20 eV at 0.6 V.s/cm^2^. The collision cell RF (Vpp) was set to 1500 V, the pre-pulse storage time was set to 12 textmus with 60 µs transfer time. The TIMS dimension was calibrated using Agilent ESI LC/MS tuning mix (*m/z*, 1/K0): (622.0289, 0.9848 Vs/cm^2^), (922.0097, 1.1895 Vs/cm^2^) and (1221.9906, 1.3820 Vs/cm^2^) in positive mode. For filtering glycopeptide-specific PASEF precursors a modified user-defined polygon filter was implemented in the acquisition based on reduced ion mobility (1/K0) and monoisotopic mass *m/z*, with the following boundaries (*m/z*, 1/K0): (613, 1.016 Vs/cm^2^), (1220, 1.451 Vs/cm^2^), (1400, 1.605 Vs/cm^2^), (1700, 1.605 Vs/cm^2^) and (1700, 0.798 Vs/cm^2^). Human plasma samples were further fragmented using a constant CE value (no linear interpolation of the CE with reduced ion mobility) starting from 40 eV to 100 eV in 7 individual runs with and without the glyco polygon PASEF method. SGP and asialo-SGP were also fragmented with constant CE values starting from 40 to 80 eV in 5 individual runs with the standard PASEF method. For human plasma glycopeptides we further implemented a stricter polygon as follows (*m/z*, 1/K0): (800, 1.05 Vs/cm^2^), (1220, 1.30 Vs/cm^2^), (1400, 1.4 Vs/cm2), (1700, 1.40 Vs/cm^2^), (1700, 1.10 Vs/cm^2^) and (800, 0.80 Vs/cm^2^). The efficiency of this specific method was tested using shorter LC gradients (see “results” for further discussion).

Plasma and neutrophil measurements were replicated on a second timsTOF Pro device at the Bruker (Bremen) lab in triplicates, using the four different methods as follows: PASEF, PASEF SCE, PASEF with glyco polygon and PASEF with SCE and glyco polygon. The samples were separated on the nanoElute (Bruker Daltonik) coupled on-line to a timsTOF Pro mass spectrometer (Bruker Daltonik). Peptides and glycopeptides were analytically separated on an Ion Optics nanoUHPLC column (75 µm x 25 cm, 1.6 µm, C18; Ion Optics, Australia), heated to 50 °C at a flow rate of 400 nL/min. LC mobile phases A and B were water with 0.1% formic acid (v/v) and ACN with formic acid 0.1% (v/v), respectively. The nanoLC was coupled to the timsTOF Pro using the CaptiveSpray (Bruker Daltonik). All the samples were separated using the same 150 min gradient as used for the previous neutrophil and plasma samples, while other parameters were kept the same for comparative analysis between the two labs apart from the TIMS region voltages that were set to Δ6 to 55V and TIMS RF to 450Vpp.

### Data Analysis

The fragmentation spectra from all precursors with charge-state >2 were extracted from the recorded Bruker .d format files and stored in Mascot Generic Format (MGF) files with the in-house developed tool HlxlToolchain. The conversion procedure consisted of two steps. In the first step, fragmentation spectra of the same precursor were combined into a single spectrum. Matching of the precursors was performed with the following tolerances: precursor *m/z* ± 20 ppm, retention time ± 60 s, and mobility ± 5 %. Spectral data in “quasi-profile” mode were extracted using the timsDATA 2.17.1.2-beta API obtained from Bruker. Combination of the spectra was achieved by summing peak intensities of all spectra across complete “quasi-profile” *m/z* grid. The final summed spectrum was generated through removal of zero intensity peaks by binning summed “quasi-profile” spectrum in *m/z* bins of 50 ppm. In the second step, each combined spectrum was deisotoped (isotopes were reduced to a single peak at m/z of charge state of 1), and TopX filtered at 20 peaks per 100 Th. Together with the conversion procedure, an MGF-meta file was automatically created that contained information on the precursor intensity, mobility (1/K0), CCS, and monoisotopic mass. The CCS values were calculated according to the Mason-Schamp equation(35, 36), with temperature being set to 305 K and the molecular weight of N2 as the TIMS gas. The MGF files were searched with another in-house tool, HlxlGlyco, that searched specifically for 8 glycan-oxonium ions (H_14_C_8_N_1_O_5_ (204.0872), H_12_C_6_O_8_P_1_ (243.0270), H_16_C_11_N_1_O_7_ (274.0927), H_18_C_11_N_1_O_8_ (292.1032), H_24_C_14_N_1_O_10_ (366.1400), H_34_ C_20_N_1_O_14_ (512.198), H_34_C_20_N_1_O_15_ (528.1928) and H_41_C_25_N_2_O_18_ (657.2354)) in the MSMS spectra to pre-select the precursors that were likely *N*-glycopeptides from the large number of spectra. Together with the search, each precursor was associated with a glycan M-score, *i*.*e*., weighted based on the intensity of the oxonium ions present in the MSMS spectra, as previously described(37). An oxonium ion meta-file was generated containing the information on precursor *m/z*, mobility, CCS and glycan M-score. The individual CE data files for the human plasma (with and without glyco polygon) were converted to MGF format and single combined MGF files were created where all spectra originating from the same precursor using: precursor m/z ± 20 ppm, retention time ± 60 s, and mobility ± 5 %, and the intensities were summed together in the final spectrum. The MGF files were searched and processed with MSFragger (version 3.4), Fragpipe (version 17.1) and Philosopher (version 4.1.0) for *N*-glycopeptides(38). Briefly, MFG files were searched against the human Uniport FASTA (UP000005640 reviewed with 20,371 entries, downloaded from Uniprot on July 30, 2021) with the glyco-N-HCD workflow. The search parameters were as follows: precursor window: lower mass was set to 400 Da, upper mass was set to 5000 Da; precursor and fragment mass tolerance: ±20 ppm; enzyme: full trypsin digestion with two maximum missed cleavages; carbamidomethylation at Cys was set as fixed modification and oxidation at Met and protein N-term acetylation were set as variable modifications. Peptide filtering at 1% false discovery rate (FDR) was applied through PeptideProphet. Default parameters for *N*-glycan analysis with glycan FDR ¡1% and glycan mass tolerance 50 ppm were used. The human neutrophil samples were first searched against the same human FASTA file, using Mascot version 2.7.0.0(39), using precursor mass tolerance ± 20 ppm and fragment mass tolerance ± 50 ppm; enzyme: semi-specific trypsin + Glu-C digestion; carbamidomethylation at Cys was set as fixed modification and oxidation at Met and protein N-term acetylation were set as variable modifications. This subsequently yield the top 500 proteins. The human neutrophil MGF files were searched in MSFragger against the top 500 protein database with the glyco-N-HCD workflow, together with semi-specific digestion (trypsin-GluC) at a maximum of two missed cleavages at Lys/Arg/Asp/Glu. The output was filtered for the *N*-glycopeptides, spectrum scan numbers of annotated spectra were merged with the glycan-oxonium result from HlxlGlyco tool.

Next to the MSFragger-based approach, the final neutrophil and plasma samples were additionally searched and processed through a combination of MSConvert and Byonic, in line with previous reports(34, 40), to provide validation of the MSFragger results. Briefly, raw files originating from the timsTOF Pro experiments were converted to MGF format using MSConvert (3.0.21328-404bcf1), with scanSumming on precursorTol=0.05, scanTimeTol=5 and IonMobilityTol=0.01. The resulting MGF files were searched with Byonic (v4.4.1), using a list of 279 *N*-glycans set as common1(34), together with fixed Cys carbamidomethylation and rare Ser/Thr/Tyr phosphorylation, Met/Trp oxidation, and peptide- and protein-N-terminal Glu/Gln pyroglutamic acid formation. Semi-specific digestion was allowed with 3 missed cleavages, at Lys/Arg for the plasma samples and Lys/Arg/Asp/Glu for the neutrophil samples. In alignment with previous studies, PSMs resulting from the Byonic searches were curated to have a score of ≥ 150 and —log prob— value of ≥ 1.5.

Further downstream analysis and visual representation of the results was performed with the R(4.03) packages extended with ggplot2 (2.3.3.5) and eulerr (6.1.1) for data visualization. For visualization of the glycan species, we followed the recommendations of the Consortium for Functional Genomics(41). Glycan cartoons were constructed and exported from GlycoWorkbench(42).

## Results

### Optimization of the Ion Mobility Region of Interest for Targeted Analysis of Glycopeptides

We first optimized PASEF data acquisition on purified single glycoproteins guided by the sensitive detection of the diagnostic glycopeptide derived oxonium ions. Oxonium ions, singly-charged mono- and oligosaccharides originating from glycopeptide fragmentation were selected as glycopeptide diagnostic species (*m/z*=204.0872 (HexNAc), *m/z*=274.0921 (NeuAc-H2O), *m/z*=292.1032 (NeuAc), *m/z*=366.1400 (HexNAc-Hex), *m/z*=528.1928 (HexNAc-Hex2), and *m/z*=657.2354 (HexNAc-Hex-NeuAc)) to provide a view on the location of the glycopeptides inside the mobilogram of all precursor ions (Figure 2). Precursor ions with any of these diagnostic ions were observed to cluster inside the ion mobility region comprised of 1/K0 = 0.8-1.4 and *m/z* = 650-1700, respectively. To distinguish between chemical noise and oxonium ion containing precursor ion signals, we calculated a weighted oxonium ions score (M-score) that allowed us to select only those MS/MS spectra that are likely originating from *N*-glycopeptides(37). As expected, most of the MS/MS spectra had an M-score < 0.5, as the sample contained also many non-modified peptides. Previously, it has been suggested that an M-score > 1.3 leads to identification of *N*-glycopeptide precursors with an FDR< 2.5 %(37). Indeed, application of this M-score cut-off led to a selection where almost all the precursors containing at least 2 oxonium ions were inside the ion mobility polygon (Figure 2A and 2B). Glycopeptide searches on the data acquired with stepped collision energy for the phosphatase TNAP (Figure 2 and Figure S1-S2) and standard PASEF method for HRG (Figure S1, S3 and S4), proteins that have complex glycosylation especially mono- and disialylated glycans (N_4_H_5_S_1_ N_4_H_5_S_2_) (Figure S1), validated that *N*-glycopeptides are separated in the ion mobility dimension from non-modified peptides (Figure 2C). To gain insight into this separation, a linear model (lm) was optimized that optimally separates non-modified and *N*-glycopeptides. This lm was used to calculate the Euclidean distances of each identification to the linear model in the mobility dimension (y-axis). The density plot of the calculated distances demonstrates that precursors generating oxonium ions (annotated glycopeptides) indeed separate from nonmodified peptides (Figure 2D), showing that the extra dimension of ion mobility helps to improve the level of detection. We additionally plotted the intensity distribution of all ions and observed a major reduction in chemical noise with no significant loss in annotated glycopeptides following the application of the M-score cut-off > 1.3, with at least 2 potential oxonium ions in the MS/MS spectra (Figure 2E and 2F).

**Figure 2.**
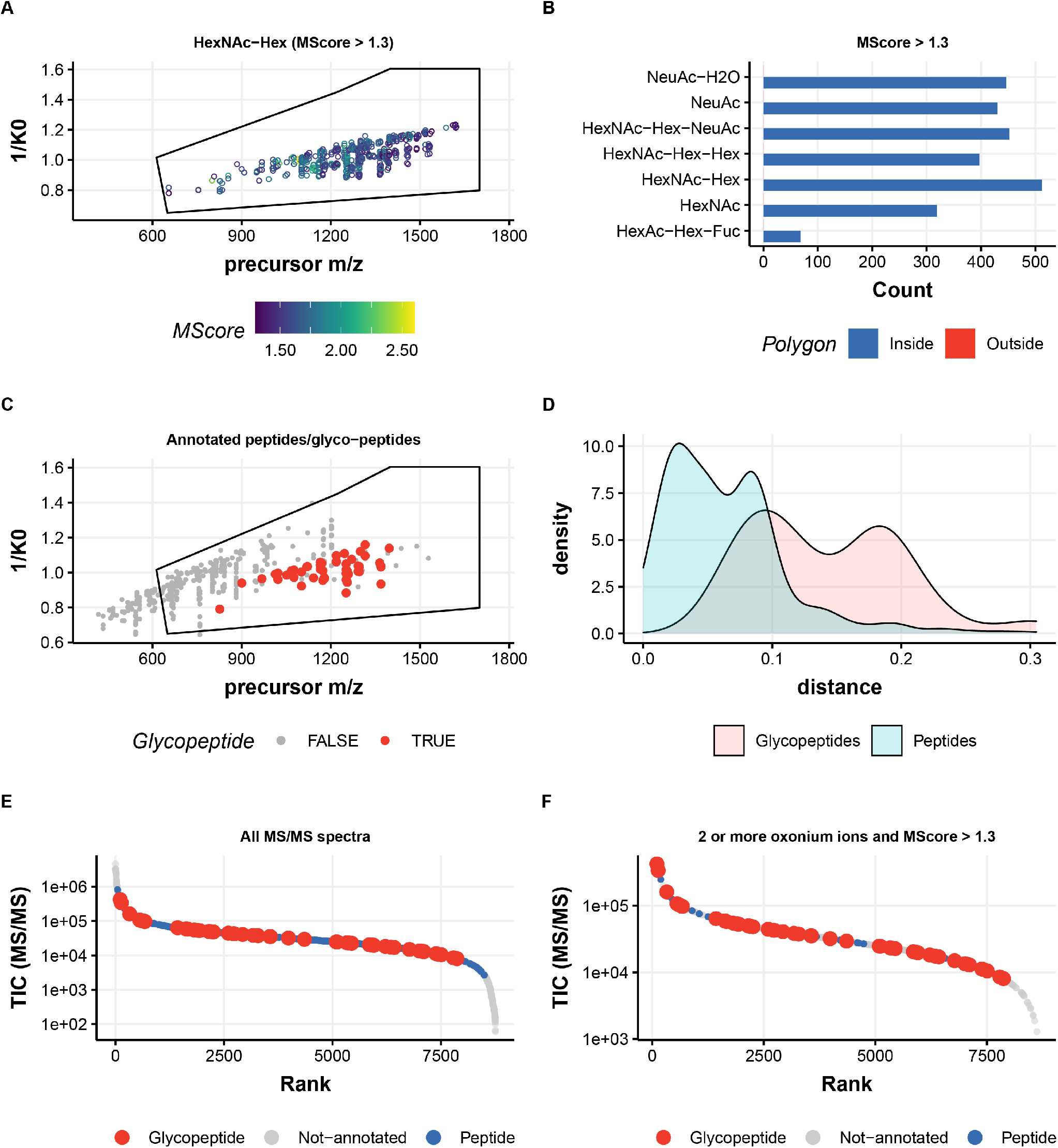
Glycopeptide identification from the purified tissue nonspecific alkaline phosphatase (TNAP) protein on timsTOF Pro. **(A)** Distribution of the precursor ion signals containing *m/z* 366.14 (HexNAc-Hex) oxonium ions, following an M-score cut-off > 1.3. **(B)** Counts of all the glycan diagnostic oxonium ions for TNAP glycopeptides demonstrate localization of all multiply charged *N*-glycopeptides precursors inside the polygon. **(C)** Distribution of the precursor ion signals in *m/z vs*. ion mobility (1/K0) for annotated peptides and *N*-glycopeptides and **(D)** density diagram displaying the physical separation of these species in the mobility space. (E) Ranked distribution of the ion signals for their intensity (TIC(MS/MS)) *vs*. rank for all classes of ions (noise in gray, annotated non-modified peptides in blue and glycopeptides in red) and (F) ranked distribution of the ion signals following the threshold of at least 2 oxonium ions and an M-score cut-off > 1.3 for the identification of glycopeptide precursor on the ion signals.

### Collision Energy Optimization

As evidently glycopeptides have different gas-phase fragmentation behaviour compared to non-modified peptides, previously optimized settings on the standard PASEF ion mobility based collision energy were not optimal to properly fragment *N*-glycopeptides on the timsTOF Pro. Low collisional energies allow resolving specific glycan structural motifs of *N*-glycopeptides, while higher collisional energies provide information of the site of glycan-protein attachment, peptide fragment ions and the assignment of features related to glycan core structures such as core-fucosylation(43, 44). Stepped collision energy (SCE) MS/MS combines these two worlds and has been widely used in the high-throughput identification of intact glycopeptides as it generates the most informative and abundant fragment ions for both glycan and peptide sequencing(15). We thus developed and optimized a stepped PASEF method on the timsTOF Pro for the identifications of the *N*-glycopeptides based on the optimal detection of oxonium ions (Figure S5 and S6) for identification of potential *N*-glycopeptides. The resulting curve (Figure S6C) is sensitive for the detection of specific glycan-oxonium ions and led to the successful identification of 28 unique *N*-glycopeptides originating from the purified protein phosphatase TNAP (Figure 2).

### Performance on more complex samples

We next subjected glycopeptides, derived from neutrophils, post-desalting, to RP-LC-TIMS-MS/MS on the timsTOF Pro with the broad and inclusive polygon and PASEF SCE fragmentation. First, we evaluated the performance of this glycoproteomic workflow in properly sequencing heterogenous *N*-glycopeptides, including sialylation, fucosylation, as well as pauci-, phospho- and high-mannose glycans that commonly occur on neutrophil glycoproteins and their resulting glycopeptides (Figure 3)(34, 40, 45, 46). *N*-Glycopeptides originating from the neutrophils (Figure 4 and Figure S7) clustered in a specific region of the ion mobility. Importantly, all high-scoring (M-score > 1.3) oxonium ion containing precursors were clustered inside the polygon (Figure S7). The glyco-oxonium ion containing precursors inside the polygon were rather indistinguishable, demonstrating that separation in the TIMS is primarily based on the intact glycopeptide *m/z* and less dependent on the exact nature of the glycan moiety (Figure S8). We could identify almost 1500 proteins with a semi-specific search at FDR <1%, examples being the abundant neutrophil glycoproteins lactotransferrin and myeloperoxidase. The identified proteome was in congruence with the results from a recent neutrophil proteomics study(47). In our optimized stepped glyco-PASEF method we identified 440 unique *N*-glycopeptides (222 present in all three replicates) from 54 glycoproteins (Table S1, Figure S9). In comparison, using a generic PASEF method (without glyco-polygon and SCE) and glyco-polygon PASEF (without SCE) we only detected 244 and 196 *N*-glycopeptides (across three replicates), from 35 and 27 glycoproteins respectively. In other words, the SCE optimized approach increased the number of identified *N*-glycopeptides across triplicate runs on average by 2.2-fold (Figure S9). When visualizing the mobility of the annotated *N*-glycopeptides versus *m/z*, there was a clear physical separation between the glycopeptides and majority of non-modified peptides (Figure 4C). Using the same lm model calculation approach as described above and calculating the Euclidean distances for the precursors with M-score > 1.3, we observed that the density of the *N*-glycopeptides from the neutrophil samples were also separated from the non-modified peptides in the ion mobility domain (Figure 4D).

**Figure 3.**
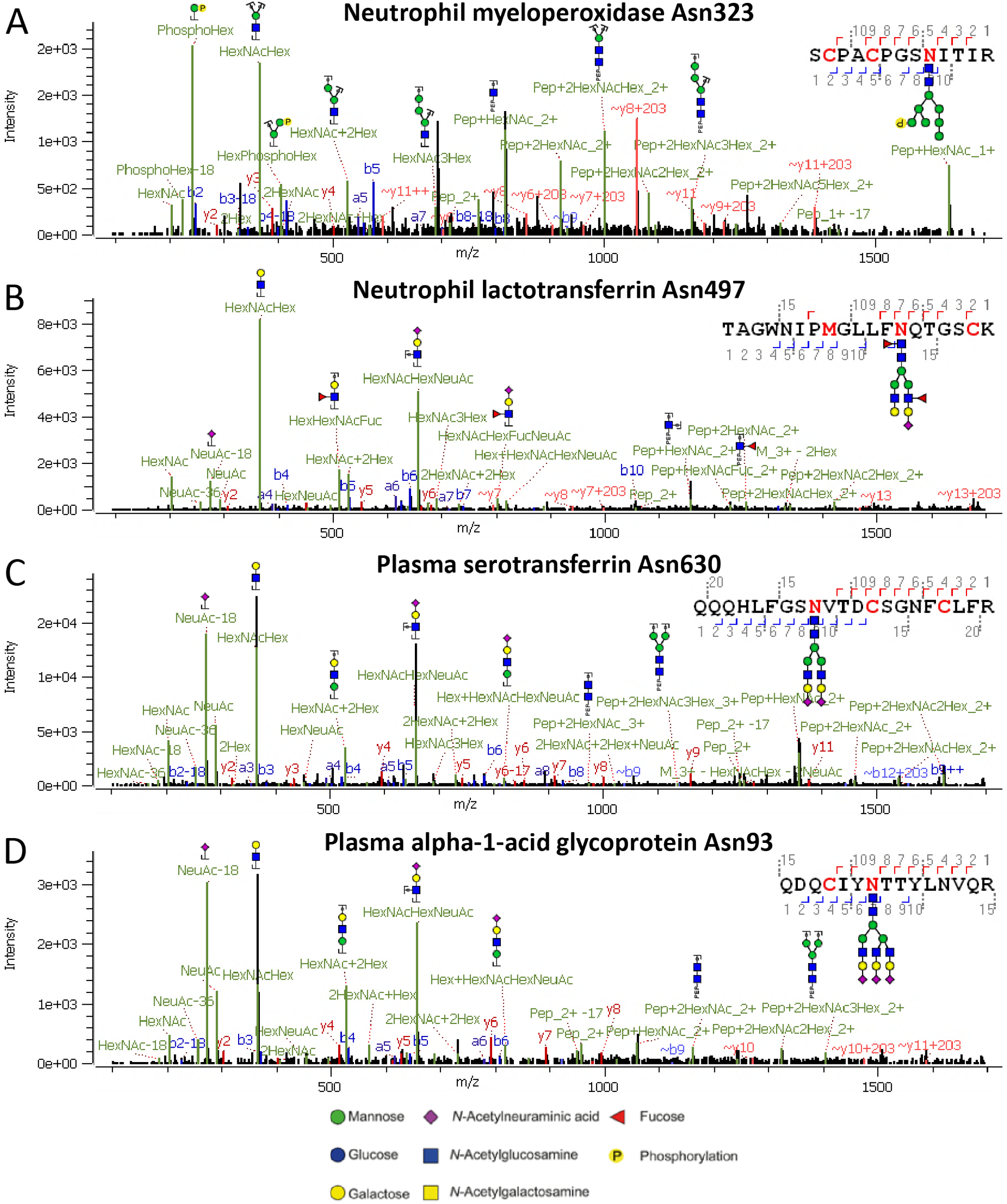
Illustrative annotated tandem mass spectra of *N*-glycopeptides, displaying the observed diverse glycosylation categories that can be identified using the timsTOF Pro. **(A)** Phoshomannose glycosylation on a neutrophil myeloperoxidase glycopeptide at Asn323, **(B)** antennary fucosylation on a neutrophil lactotransferrin glycopeptide at Asn497, **(C)** sialylation on a glycopeptide from plasma serotransferrin at site Asn630 and **(D)** triantennary species on a glycopeptide from plasma alpha-1-acid glycoprotein at site Asn93. These spectra were obtained by summation of spectra acquired at SCE collision energies. These tandem mass spectra demonstrate the performance of the stepped SCE-MS/MS fragmentation on the timsTOF Pro resulting in glyco-oxonium ions (approximately *m/z* 200-700), peptide backbone fragments (b- and y-ions) and glycan residue losses (B- and Y-fragments). Glycan nomenclature used in glycopeptide definitions are delineated at the bottom of the figure.

**Figure 4.**
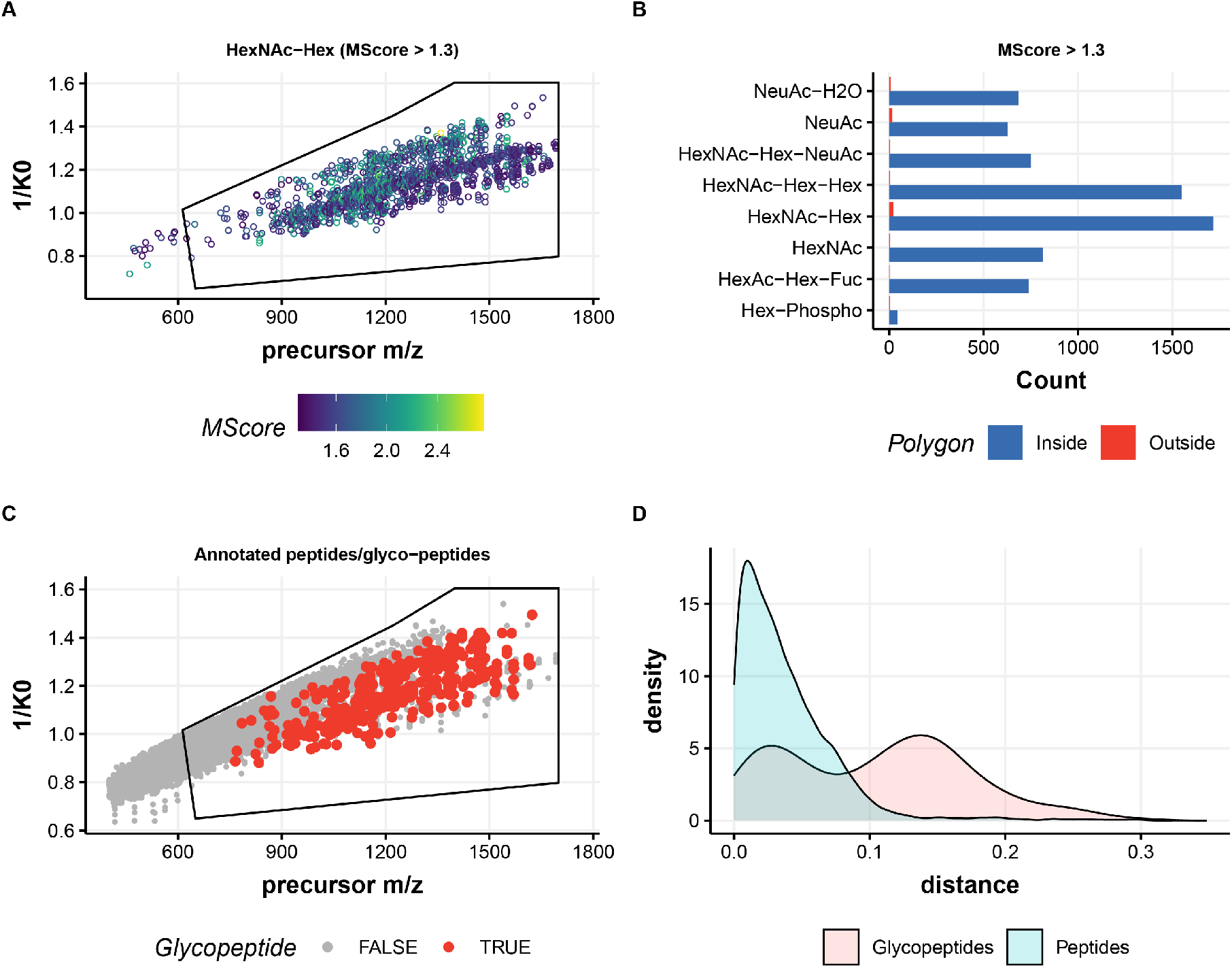
Identification of *N*-glycopeptides originating from in human neutrophils. **(A)** Distribution of the precursor ion signals containing *m/z* 366.14 (HexNAc-Hex) oxonium ions, following an M-score cut-off > 1.3. **(B)** Counts of all the glycan diagnostic oxonium ions for neutrophil glycopeptides demonstrate localization of nearly all multiply charged *N*-glycopeptides precursors inside the polygon. **(C)** Distribution of the precursor ion signals in *m/z vs* ion mobility (1/K0) for annotated peptides and *N*-glycopeptides and **(D)** density diagram displaying the physical separation of these species in the mobility space.

We additionally subjected human plasma to our optimized workflow, using trypsin to digest the proteins. We were able to detect and sequence 518 unique plasma *N*-glycopeptides (275 annotated in all three replicates) originating from 81 unique glycoproteins (Table S2, Figure S11). In comparison, PASEF and polygon-PASEF (without SCE) methods could identify only 76 and 72 unique *N*-glycopeptides, respectively, from 32 and 31 glycoproteins (Table S2, Figure S10). This represents a 6.8-7.1-fold increase in the identification rate of *N*-glycopeptides when using SCE methods. Of note, SCE-PASEF (without the specific glyco-polygon) performed equally well as stepped glyco-PASEF, where we could identify 526 unique *N*-glycopeptides (288 annotated in all three replicates) from 83 glycoproteins. A total of 67.4 % of *N*-glycopeptides overlapped between these two methods with more than 90 % overlap at glycoprotein level (Figure S12). The lack of benefit is explained by the liberal polygon used to capture all glyco-peptides.

The results from both human neutrophil and human plasma samples, additionally indicate that to fully exploit the benefits of the glyco-polygon concept it has to be optimised for specific sample type. Additionally, because of the high timsTOF Pro data acquisition speed it would be possible to use more comprehensive fragmentation methods. For example, as an alternative to SCE method with two predefined CE gradients each precursor can be measured at 5 or more different CE to obtain better fragmentation patterns of both the peptide and glycan fragments. As a proof of concept, we collected human plasma data using the standard PASEF method at 7 different CE (40, 50, 60, 70, 80, 90 and 100) with and without glyco-polygon defined. The results (Figure 5, Figure S13, Figure S14 and Table S3) demonstrate a clear increase in the numbers of annotated glycopeptides, glycan M-score values and peptide ion coverage (increase in MSFragger Hyperscore) in search results where spectra acquired at different CE’s are merged into single spectrum. Additionally, using the glyco-polygon for data acquisition effectively increased the number annotated *N*-glycopeptides (545 in CE merge, polygon) by almost 12% in comparison to PASEF method without polygon (478 in CE merged). From these results it is clear that future developments of the timsTOF Pro methods for glycoproteomics should be aimed towards developing MS/MS methods allowing dynamic application of multiple collision energies. To furthermore demonstrate applicability of glyco ion selection polygon we next focussed on shorter chromatography gradients as described in the next paragraph.

**Figure 5.**
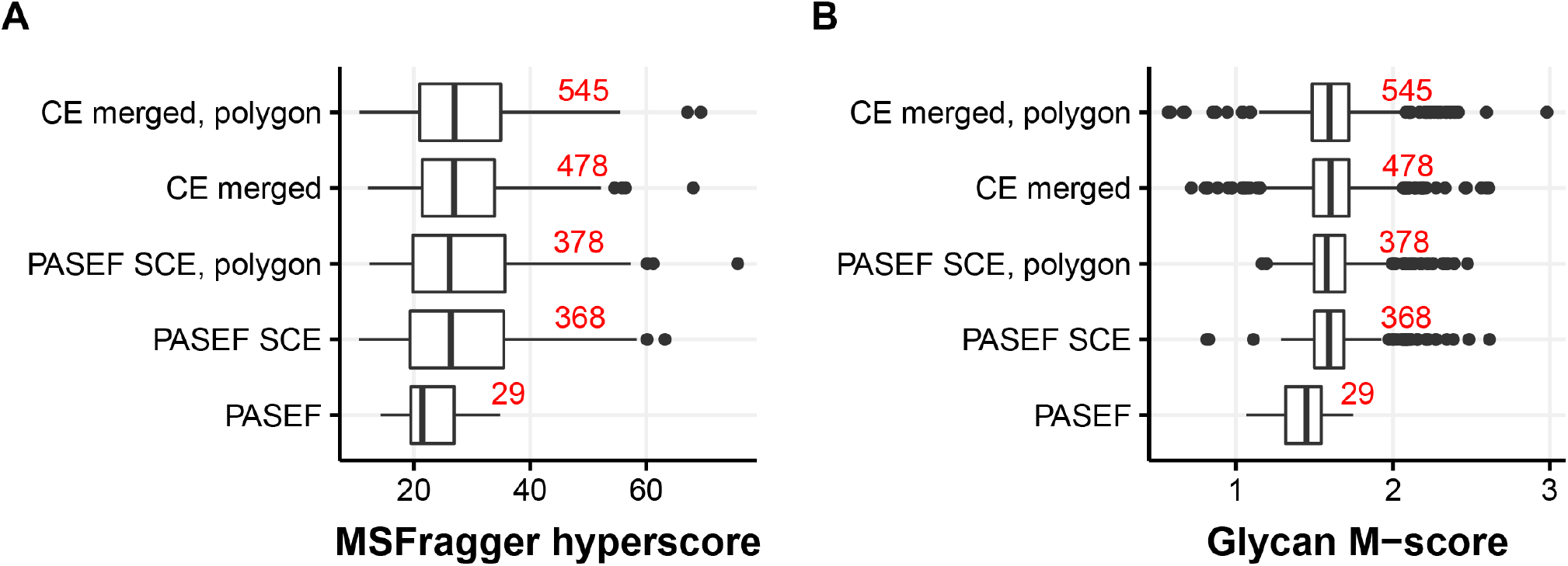
Performance or glycopeptide annotation using data acquired using PASEF, PASEF SCE and PASEF SCE glyco-polygon methods in comparison to a data set with merged CE spectra. Synthetic data files are constructed from data files collected at 7 different CE’s (40, 50, 60, 70, 80. 90 and 100) measure with (CE merged, polygon) and without (CE merged) glyco-polygon. Numbers in red represent count of unique annotated glycopeptides. **(A)** Clear increase in peptide annotation score from MSFragger can be observed in SCE data and CE merged results. **(B)** Application of different CE values significantly improve glycan score of MS/MS spectrum.

### Focussing leads to increased analytical depth

Having ascertained that the glyco-oxonium ion containing precursors cluster in a specific ROI, we built a stricter glycopeptide polygon (based on the MSFragger annotations of the glyco PSMs from the broad inclusive glyco-polygon SCE PASEF results) comprised of 1/K0 1.5 to 1.4 for *m/z* 800 to 1700, respectively (Figure 6, Figure S15), to include only high-scoring and confident *N*-glycopeptides to investigate if there was any advantage of the glyco-specific ROI in ion mobility as well as its fast performance compared to SCE PASEF method. Part of the setup includes an increase in TIMS gradients that compress the ion current in narrower LC elution peaks, thereby providing higher ion counts in individual MS/MS scans. The sensitivity and the efficiency of the method was tested using sequentially shorter gradient runs on human plasma sample. For the same plasma sample, we identified 452 unique *N*-glycopeptides (mean across three replicates) from 74 glycoproteins using the polygon method compared to 376 unique *N*-glycopeptides from 67 proteins using the non-polygon method on a 90-minute gradient (Figure 6E-F). As expected, the new method retained better performance in subsequently shorter gradients as well (Figure 6E-F), the largest difference presenting itself at a 30 min gradient with the detection of approximately 1.5-fold more unique *N*-glycopeptides peptides when the strict polygon was used. As the complexity and the dynamic range of mass spectrometers are expected to increase further in the coming years, this indicates that the polygon (*i*.*e*., focussed) method will provide superior performance.

**Figure 6.**
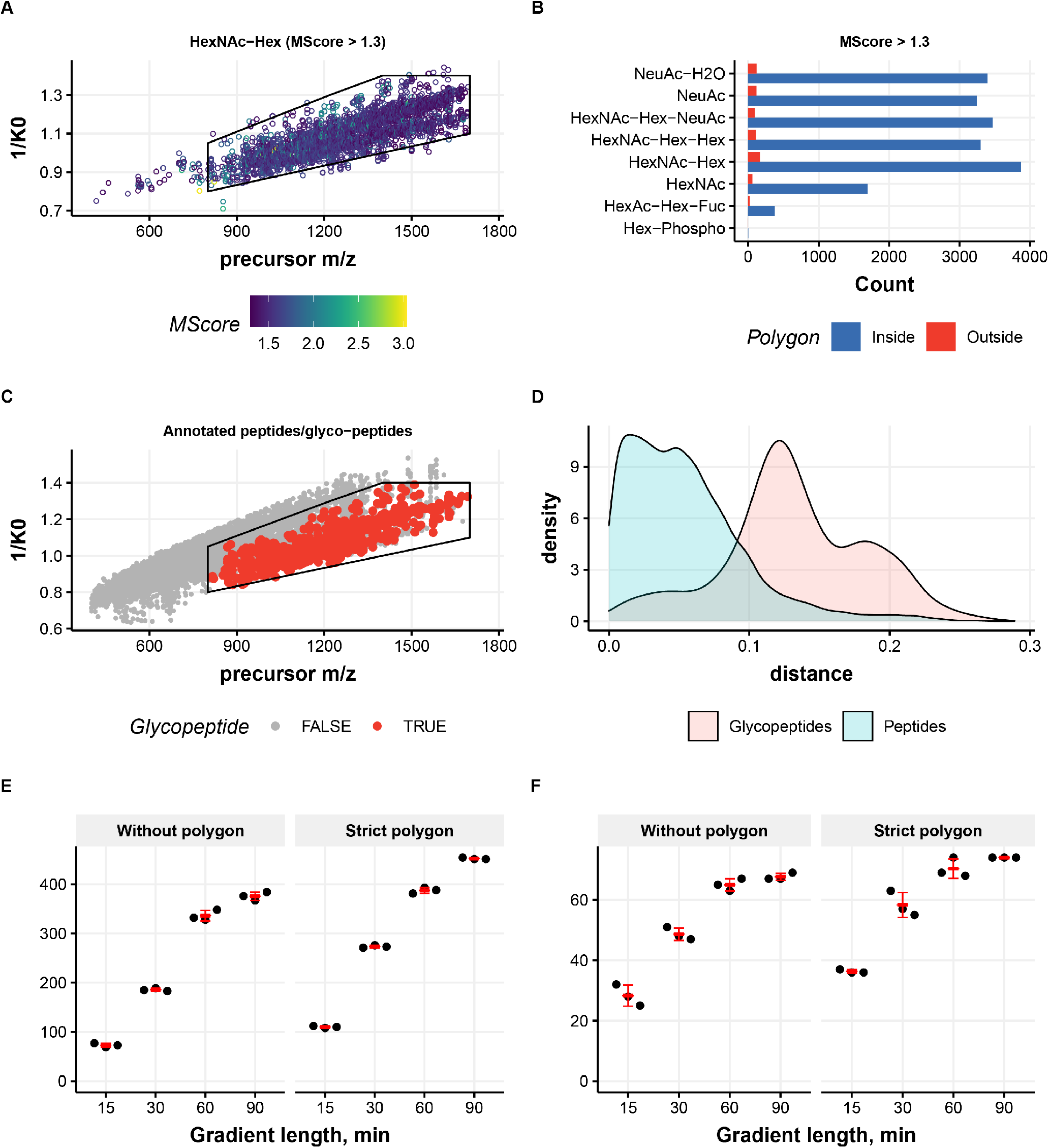
Identification of *N*-glycopeptides originating from human plasma and high-throughput glycoproteomics. **(A)** Distribution of the precursor ion signals containing *m/z* 366.14 (HexNAc-Hex) oxonium ions, following a M-score cut-off >1.3. **(B)** Counts of all the glycan diagnostic oxonium ions for plasma glycopeptides demonstrate localization of all multiply charged *N*-glycopeptides precursors inside the stricter polygon. **(C)** Distribution of the precursor ion signals in *m/z vs*. ion mobility (1/K0) for annotated peptides and *N*-glycopeptides demonstrate this smaller polygon contains most of the *N*-glycopeptides and anything outside this box can be ignored (noisy MS/MS spectra). **(D)** Density diagram displaying the physical separation of the non-modified peptides and *N*-glycopeptides in the mobility space is better with this smaller polygon. **(E)** Unique glycopeptide and **(F)** glycoprotein detection, comparing the more strict polygon with the non glyco-specific selection (without polygon) for different gradient lengths (15 min, 30 min, 60 min and 90 min).

### Quantification of glycopeptides

Finally, we qualitatively compared the peptide glycoforms from these two complex biological samples (Figure 7). For the plasma glycoproteome we observed that several *N*-glycan compositions dominated, fully in line with previous reports (48). The glycan repertoires included di- and triantennary glycan species with varying degrees of sialylation that largely originate from liver-produced acute phase proteins such as haptoglobin, α-2-HS-glycoprotein and α-1-acid glycoprotein, partially galactosylated glycans that are mainly found on the varying subclasses of immunoglobulin G (IgG), as well as high-mannose glycans stemming from proteins like immunoglobulin M (IgM), apolipoprotein B-100 and complement C3(48). The neutrophil samples, on the other hand, distinctly showed phospho- and paucimannose glycans (and smaller) occurring on azurophilic granule proteins like myeloperoxidase, proteinase 3 and cathepsin G, highly fucosylated complex glycans on, *e*.*g*., lactotransferrin and neutrophil gelatinase-associated lipocalin, as well as high-mannose species on membrane-anchored proteins like integrin alpha-M and integrin beta-2. Again, these detections were highly consistent with what was previously reported for the same sample type, yet with different MS instrumentation(34).

**Figure 7.**
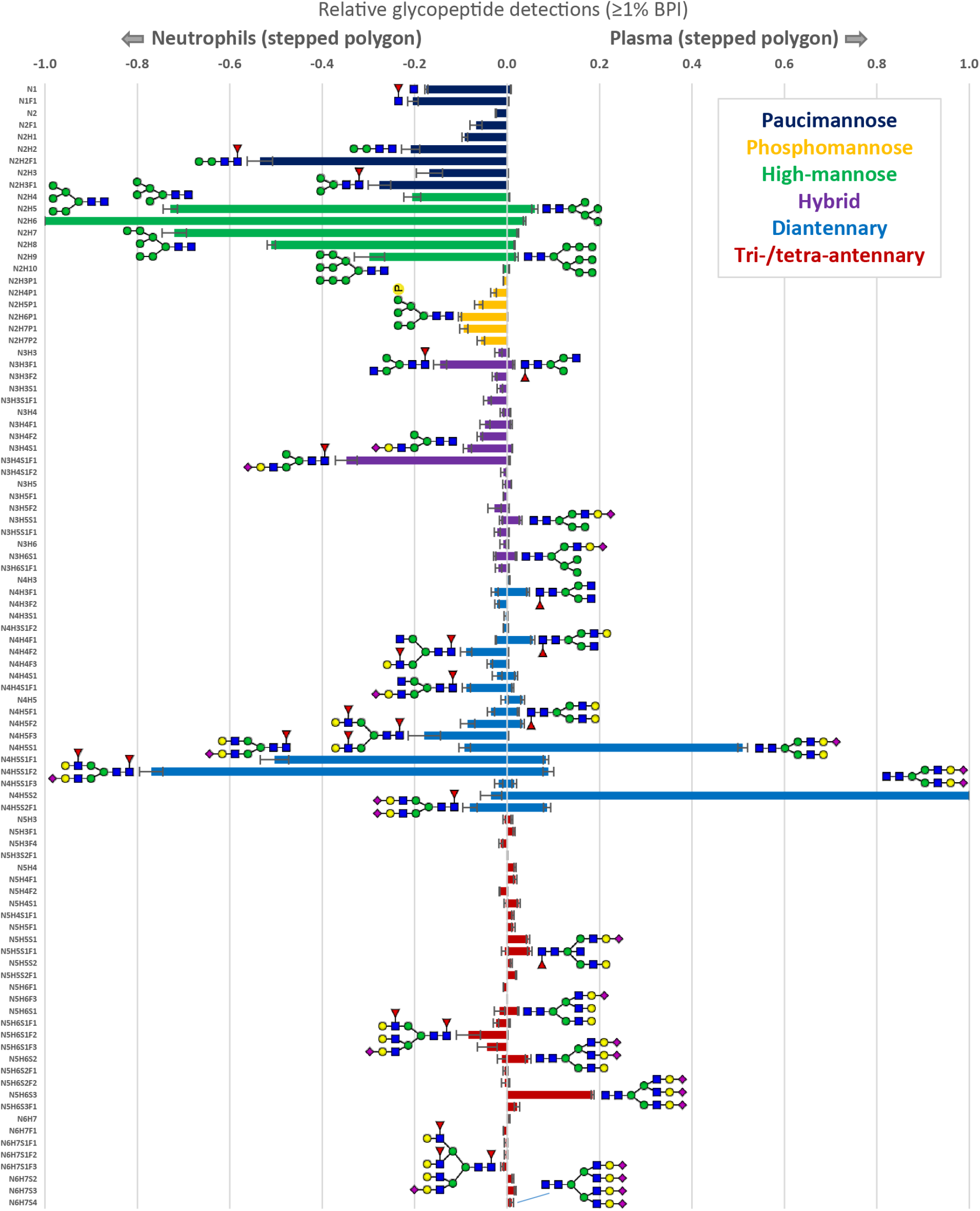
Qualitative comparison of peptide glycan repertoires observed in human neutrophils (left) and human plasma (right). Error bars represent the standard deviation for the relative quantification across triplicate injections. We assigned global glycosylation characteristics on the glycopeptides based on the monosaccharide composition. Furthermore, for visualization purposes we defined the following glycosylation characteristics: 1) paucimannose (HexNAc < 3 and Hex < 4), 2) phosphomannose (Phospho > 0), 3) high-mannose (HexNAc = 2 and Hex > 3), 4) hybrid/asymmetric (HexNAc = 3), 5) diantennary (HexNAc = 4) and 7) extended (HexNAc > 4)(34).

### Interlaboratory comparison

Proteomics and, possibly even more so, glycoproteomics experiments are often hampered by limited reproducibility, especially when comparing data obtained between different laboratories with different workflows for data acquisition and analysis.(49, 50) To test the robustness of the method presented here, we transferred the samples and methods to a second laboratory, with independent operators, but using the here-optimized method (SCE-PASEF with glyco-polygon). We overserved approximately 60 % overlap for the reproducibly detected *N*-glycopeptides (present in all three replicates) in neutrophil and plasma samples between the two laboratories (Figure S16 + Tables S1 and S2). When analysing the data along with the number and identity of the unique glycopeptides detected were very much alike, as illustrated by the nearly indistinguishable distributions of glycan moieties detected (Figure 8).

**Figure 8.**
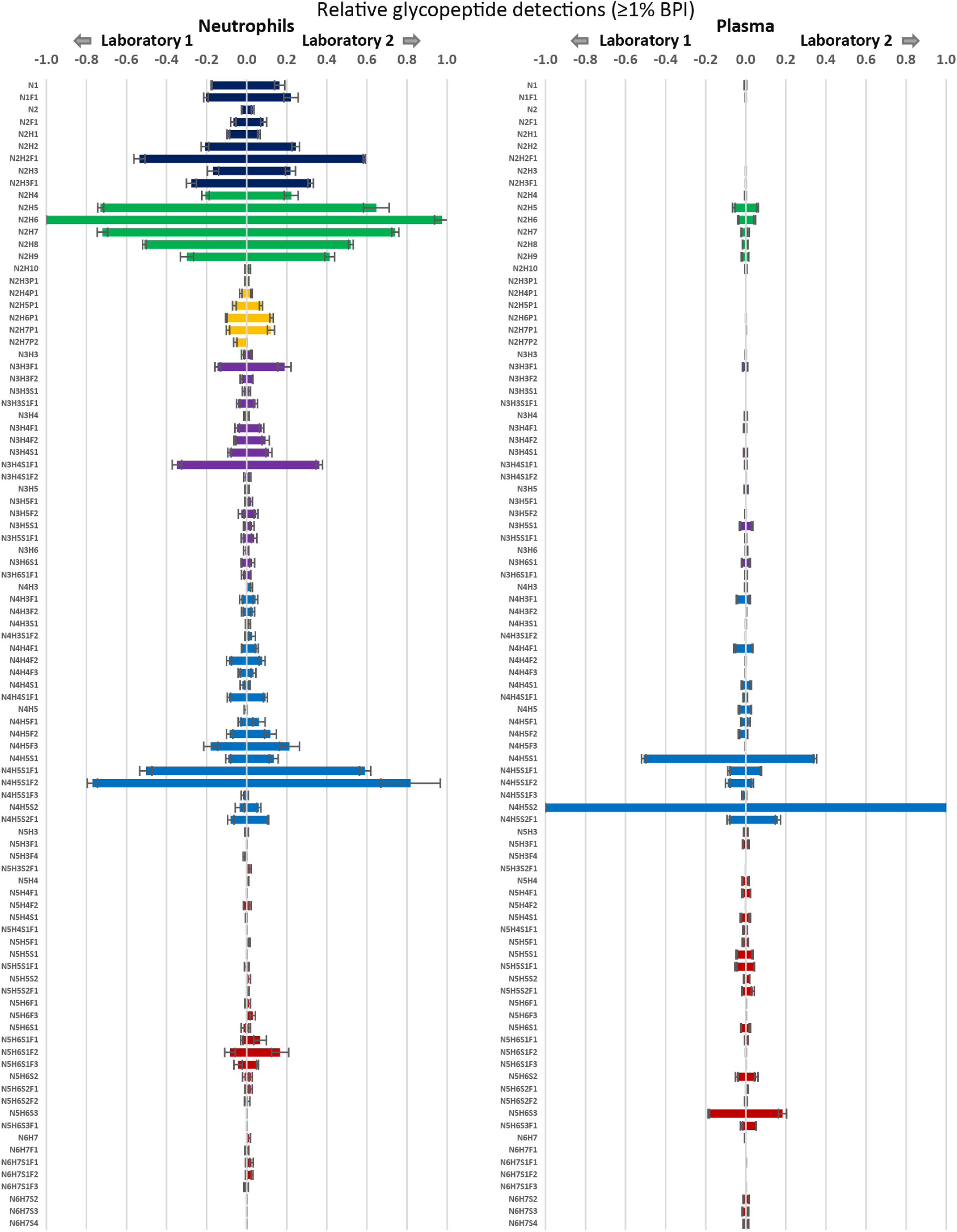
Interlaboratory comparison of the detected peptide glycan repertoires observed in human neutrophils (left) and human plasma (right). Error bars represent the standard deviation for the relative quantification across triplicate injections. Using the same samples and optimized workflow laboratory 1 and 2 respectively detected 321 and 281 glycopeptides in neutrophils and 389 and 452 glycopeptides in plasma.

## Discussion

In this study we used a nanoflow glycoproteomics workflow utilizing the advantages of both the TIMS and the PASEF methods on the timsTOF Pro. Our results indicate that physical separation can be achieved for *N*-glycopeptides in the TIMS compared to non-modified peptides, both when starting from purified single glycoproteins as well as more complex and diverse samples such as plasma. This separation helps to increase the analytical depth, which will be beneficial for future glycoproteomic analyses. A dedicated, glycan-specific polygon in the PASEF mode, together with SCE, significantly enhanced the *N*-glycopeptide identification by effectively increasing spectrum quality and maximizing the time spent on specific analytes of interest. This resulted in a boost in the identifications of *N*-glycopeptides in all samples studied, but especially so for shorter gradients. We could identify >300 unique *N*-glycopeptides from human neutrophils and >400 unique *N*-glycopeptides from plasma, resulting in a 2.2- and 7-fold increase, respectively, compared to the standard PASEF method (150 min RT gradient). Recently, 352 unique *N*-glycopeptides (89 glycoproteins)(37) have been identified in plasma (un-depleted), which was comparable with our SEC-PASEF results (452 *N*-glycopeptides, 74 glycoproteins). Of note, our merged CE with glyco-polygon resulted in >560 *N*-glycopeptides demonstrating better performance compared to affinity based glycoproteomic workflows (478 *N*-glycopeptides) on human plasma(51). Excitingly, when reducing the gradient length, the performance of the new method remains more constant, enabling either faster analysis through shorter gradients or increased analytical depth for longer gradients. Especially the first makes our workflow very attractive for glycoprotein biomarker diagnostics when larger cohorts are assessed.

However, our method also has still some drawbacks. The oxonium ions that are typically used to differentiate glycan isomers on other types of mass spectrometers fall outside the lower mass range of our experiments, effectively preventing detection of anything smaller than a HexNAc (approximately *m/z*=204). Extending the mass range towards lower *m/z* values to include HexNAc fragments and hexose oxonium ions would not only help detections in general but would also present the opportunity to distinguish some glycan isomerism (GlcNAc *vs*. GalNAc)(52, 53). An attractive strategy for the future would be to also be able to combine glycan structure or isomer detection using ion mobility separations.

The glycosylation characteristics we ultimately observed for the neutrophil and plasma samples proved to be highly congruous with earlier reports employing different instrumentation and methods(34, 48, 54). Neutrophil digests are especially challenging due to the high abundance of very small glycan species (paucimannose and smaller), labile phosphomannose residues, as well as large glycans with a complex pattern of sialylation and fucosylation on their glycopeptides(34, 40, 45, 46). Nevertheless, all of these glycosylation characteristics proved recoverable within our experiments, and while running the timsTOF Pro with standard PASEF led to a noticeable undersampling of the more complex glycans, the detection of glycopeptides across the full complexity space was allowed by the application of SCE and polygon-selection. Interestingly, in the comparison between neutrophils and plasma it was noted that sialylation (high in plasma) and fucosylation (high in neutrophils) were remarkably well-assigned according to literature expectations, even while using the same search parameters. The Fuc2 and Sia1 distinction is a pervasive analytical challenge in mass spectrometry, as these only differ by 1 Da, and are therefore easily co-isolated for fragmentation and/or misassigned in data analysis pipelines.

While it is still challenging for any MS method to ascertain what the unbiased - “true” - glycosylation profile is of any complex samples, we here report a powerful new means for glycoproteomics that is rapid, broad, and deep. We envision the use of ion-mobility assisted glycoproteomics for future clinical cohort studies and biomarker development, as well as for rapid clinical screening to achieve patient stratification.

## Acknowledgments

R.A.S. and A.J.R.H acknowledge funding through the European Union Horizon 2020 program INFRAIA project Epic-XS (Project 823839). Additionally, R.A.S. and A.J.R.H. acknowledge that this work is part of the research programme NWO TA with project number 741.018.201, which is partly financed by the Dutch Research Council (NWO). K.R.R. further acknowledges support from the NWO Veni project with project number VI.Veni.192.058.

## Contributions

SM performed experiments, data analysis and wrote the manuscript. AJ wrote software, and performed data analysis, and wrote the manuscript. FB performed experiments. ML performed experiments. YZ prepared samples, GP helped with the interpretation of the data and offered guidance for further experiments. AJRH initiated the concept and helped with interpretation of the data and wrote the manuscript. RAS designed the experiments, helped with interpretation of the data, and wrote the manuscript. KR performed data analysis, designed the experiments, helped with interpretation of the data, and wrote the manuscript.

## Data availability

The mass spectrometry proteomics data have been deposited to the ProteomeXchange Consortium via the PRIDE(55) partner repository with the dataset identifier <u>PXD034845.</u>

