## Supplementary material for "Oxonium Ion-Guided Ion Mobility-Assisted Glycoproteomics on the timsTOF Pro"

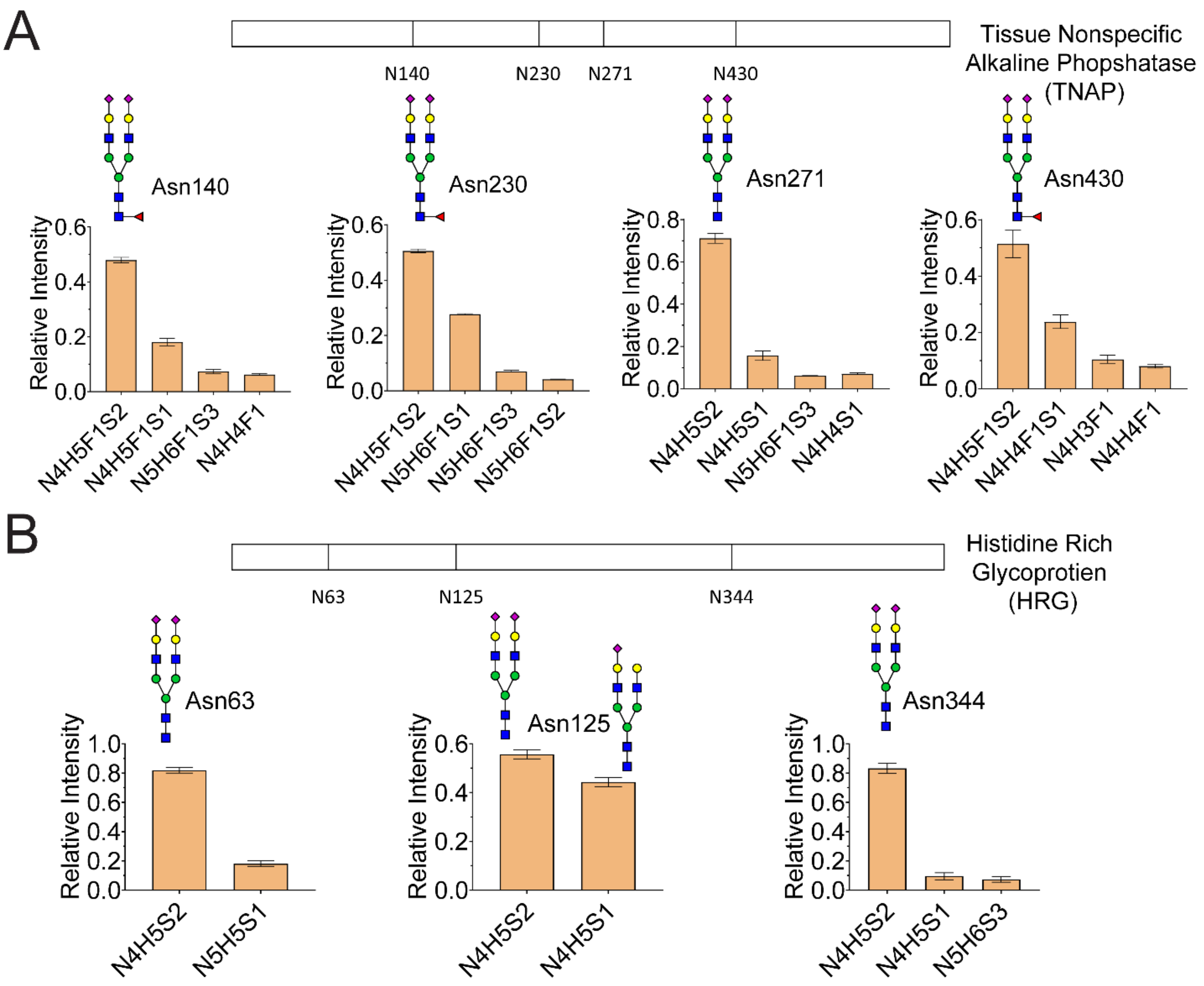


**Figure S1.** Glycan heterogeneity observed in the **(A)** 4 glycosites in the tissue nonspecific alkaline phosphatase (TNAP) and **(B)** 3 glycosites in the histidine rich glycoprotein (HRG) from human plasma. Skyline v21.1.0.146 was used to process the glycopeptides using the top three isotopic peaks from of precursor charge state +2 and +3. Integrated peak areas of each glycopeptide were summed up to represent the abundance and was used to estimate the relative abundances.

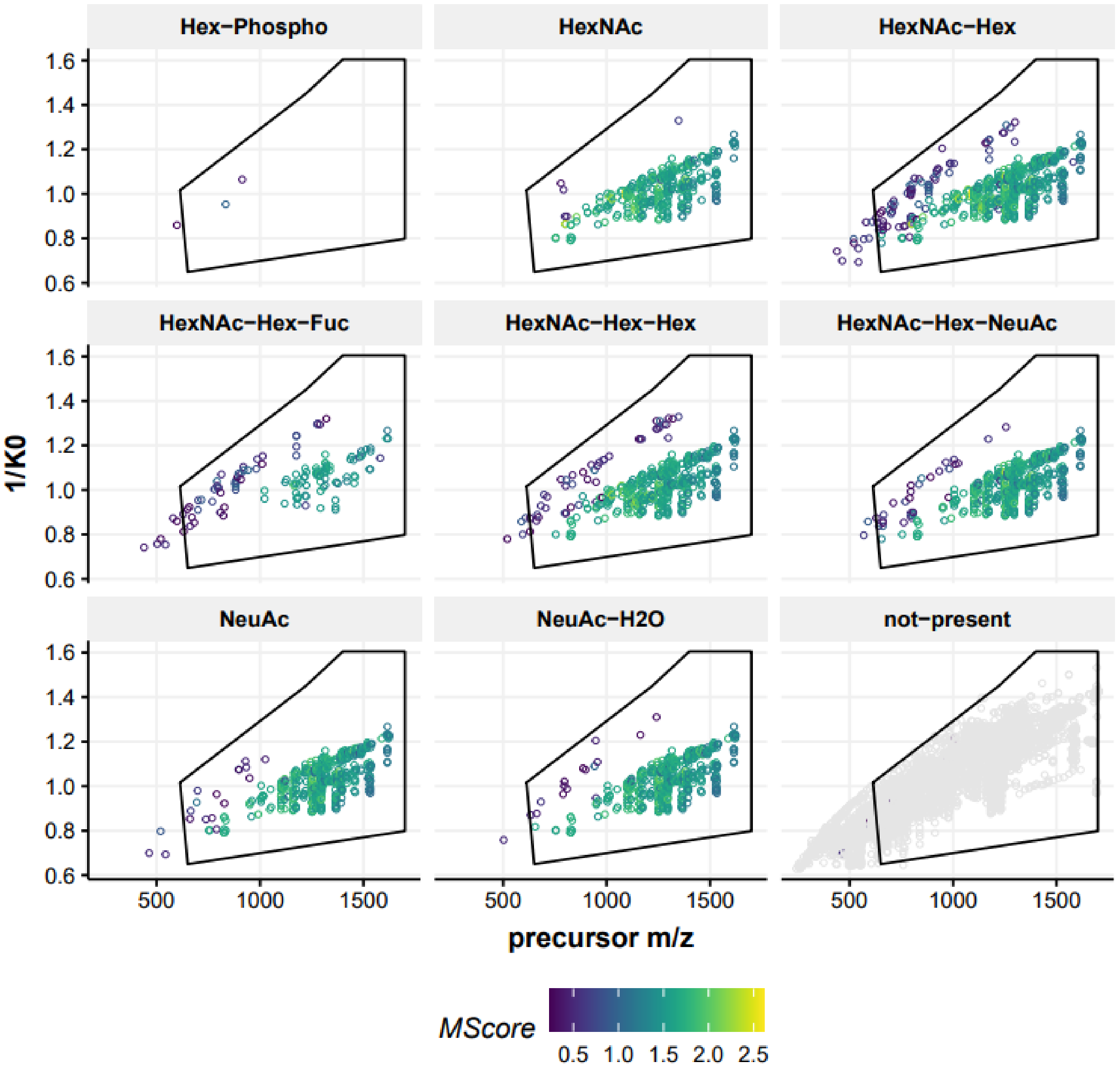


**Figure S2.** Density plot for the distribution for *m/z* vs reduced ion mobility (1/$k_0$) of all the precursor ions containing the glyco-oxonium ions 243. 0270 (Hex-Phospho), 204.0872 (HexNAc), 366.1400 (HexNAc-Hex), 512.198 (HexNAc-Hex-Fuc), 528.198 (HexNAc-Hex-Hex), 657.2354 (HexNAc-Hex-NeuAc), 292.1032 (NeuAc) and 274.0921 (NeuAc-$H_2O$) as observed for the TNAP protein. All glycopeptide precursors with M-score > 0.5 can be contained within ion selection polygon.

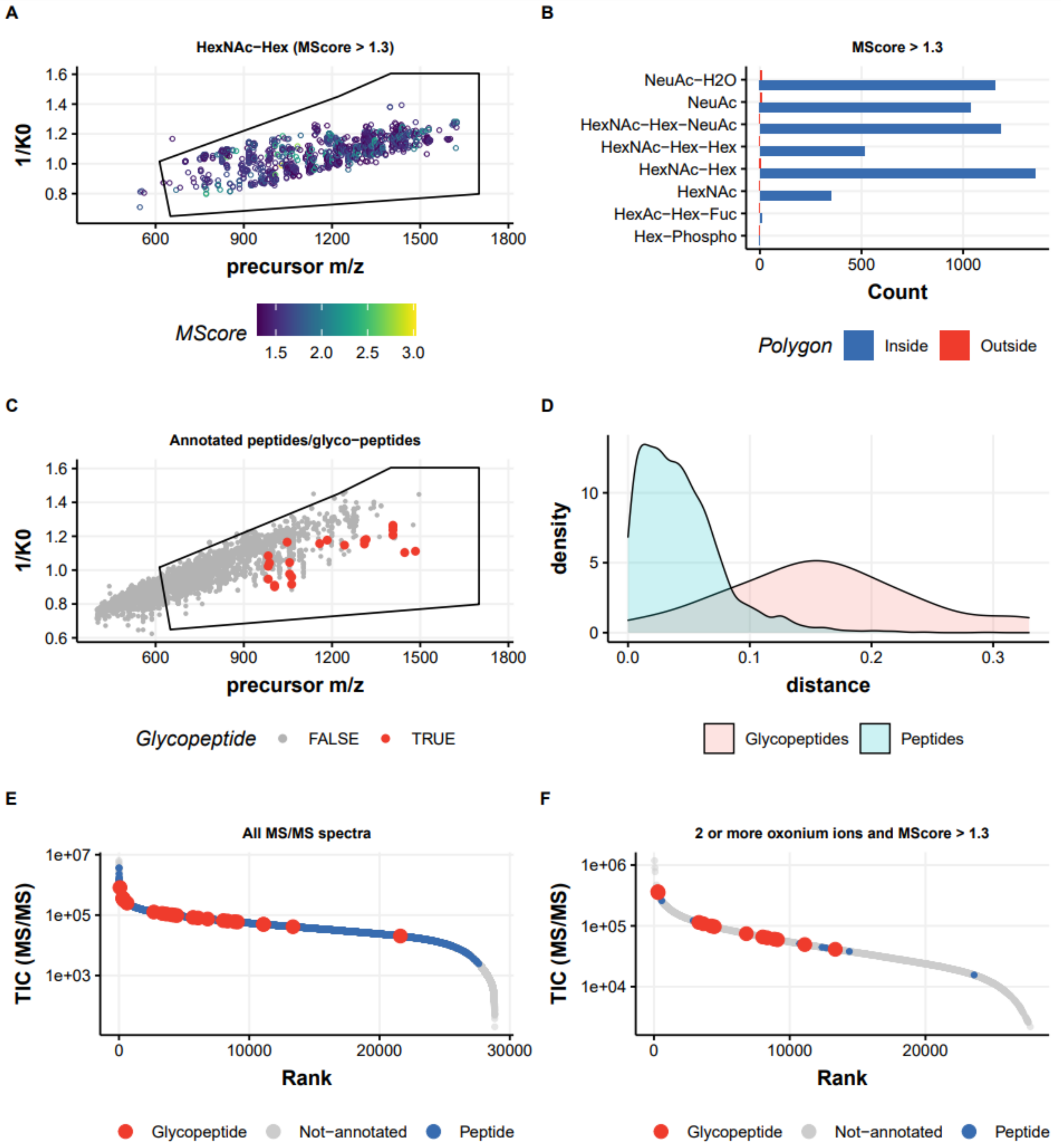


**Figure S3.** Glycopeptide identification from the plasma purified histidine rich glycoprotein (HRG) on the timsTOF Pro. Similar to Figure 2, **(A)** Distribution of the precursor ion signals containing *m/z* 366.14 (HexNAc-Hex) oxonium ions, with a MScore cut-off > 1.3. **(B)** Counts of all the glycan diagnostic oxonium ions for HRG glycopeptides demonstrate localization of all multiply charged *N*-glycopeptides precursors inside the polygon. **(C)** Distribution of the precursor ion signals in *m/z* vs ion mobility ($1/K_0$) for annotated peptides and *N*-glycopeptides and **(D)** Physical separation of these species in the mobility space. **(E)** Ranked distribution of the ion signals for their intensity (TIC(MS/MS) vs Rank for all classes of ions (noise, annotated peptides and glycopeptides) and **(F)** Same distribution following the application of 2 or more oxonium ions and MScore cut-off > 1.3 for identification of glycopeptide precursor on the ion signals. The legend for the color-coding is provided at the bottom of the figure.

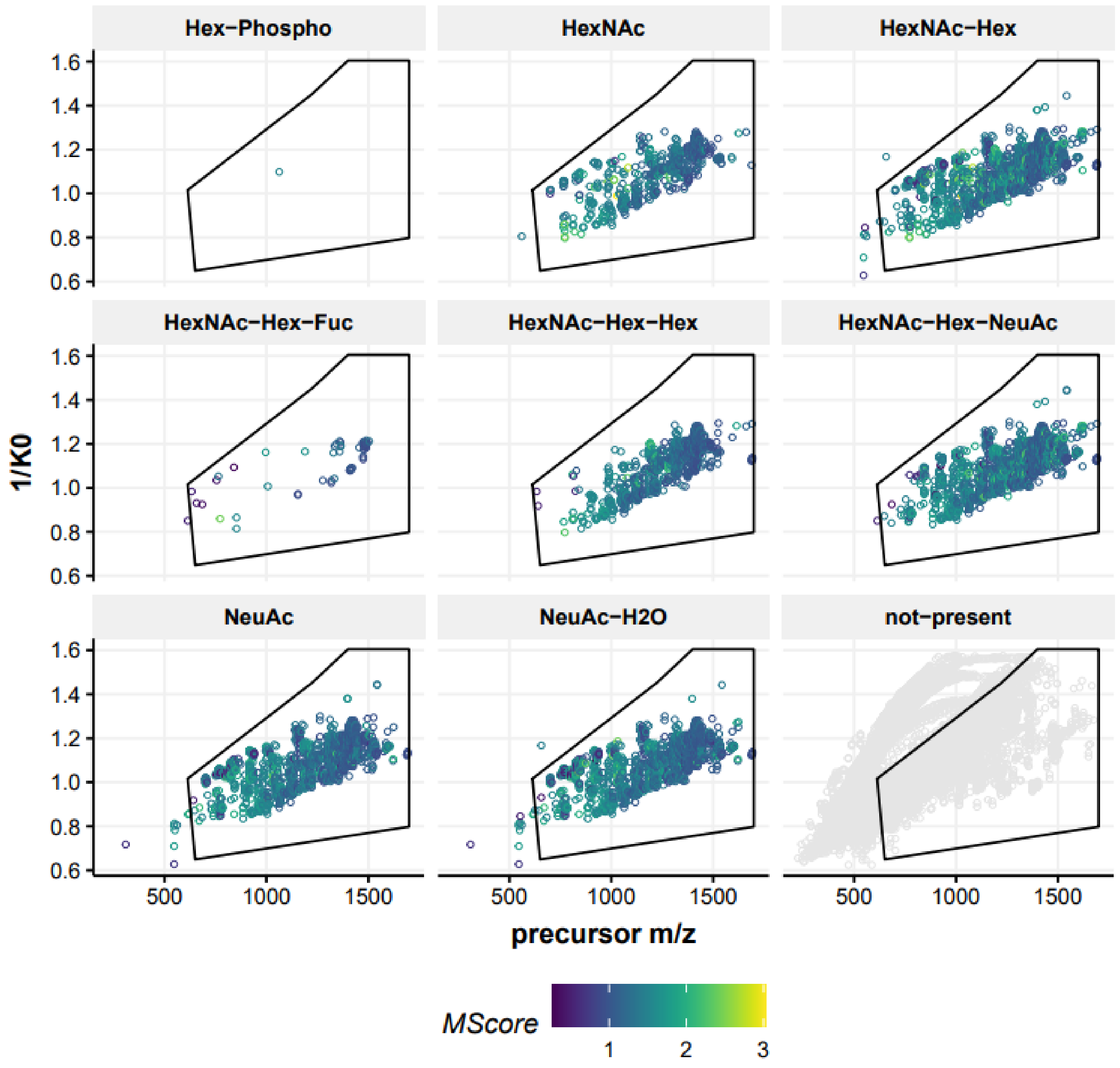


**Figure S4.** Density plot for the distribution for *m/z* vs reduced ion mobility ($1/k_0$) of all the precursor ions containing the glyco-oxonium ions 243. 0270 (Hex-Phospho), 204.0872 (HexNAc), 366.1400 (HexNAc-Hex), 512.198 (HexNAc-Hex-Fuc), 528.198 (HexNAc-Hex-Hex), 657.2354 (HexNAc-Hex-NeuAc), 292.1032 (NeuAc) and 274.0921 (NeuAc-$H_2O$) for the plasma purified HRG protein. The black polygon contains the glycopeptide precursor with M-score > 1.3.

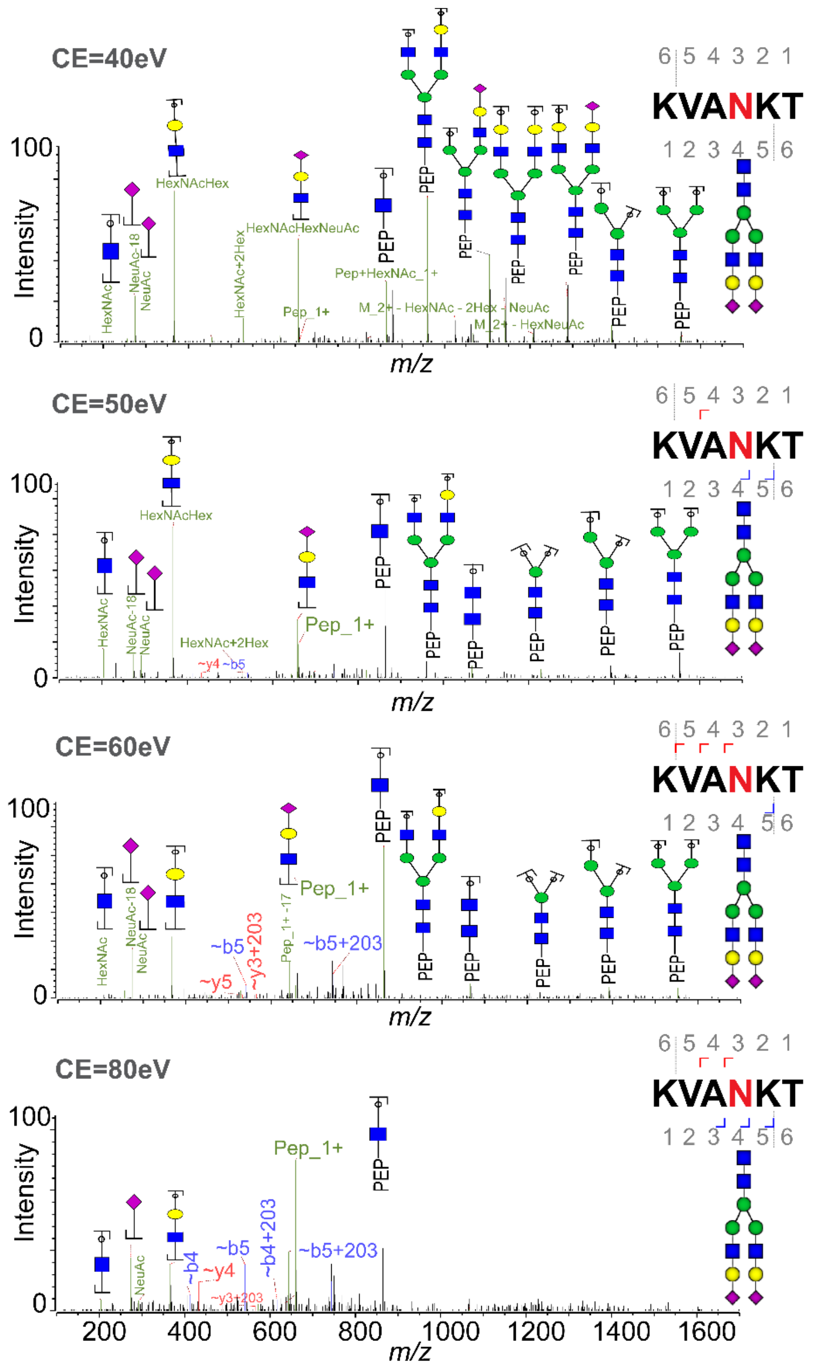


**Figure S5.** Fragmentation spectra for the SGP (α2,6-SGP) at charge state 3 at different collision energies on timsTOF Pro. At low collision energies, Y ions (glycan neutral loss) predominates, while peptide ions (b and y ions) start to appear at elevated energies. At energies higher than the optimal, glycopeptide containing ions start to disappear due to overfragmentation along with the loss of diagnostic oxonium ions.

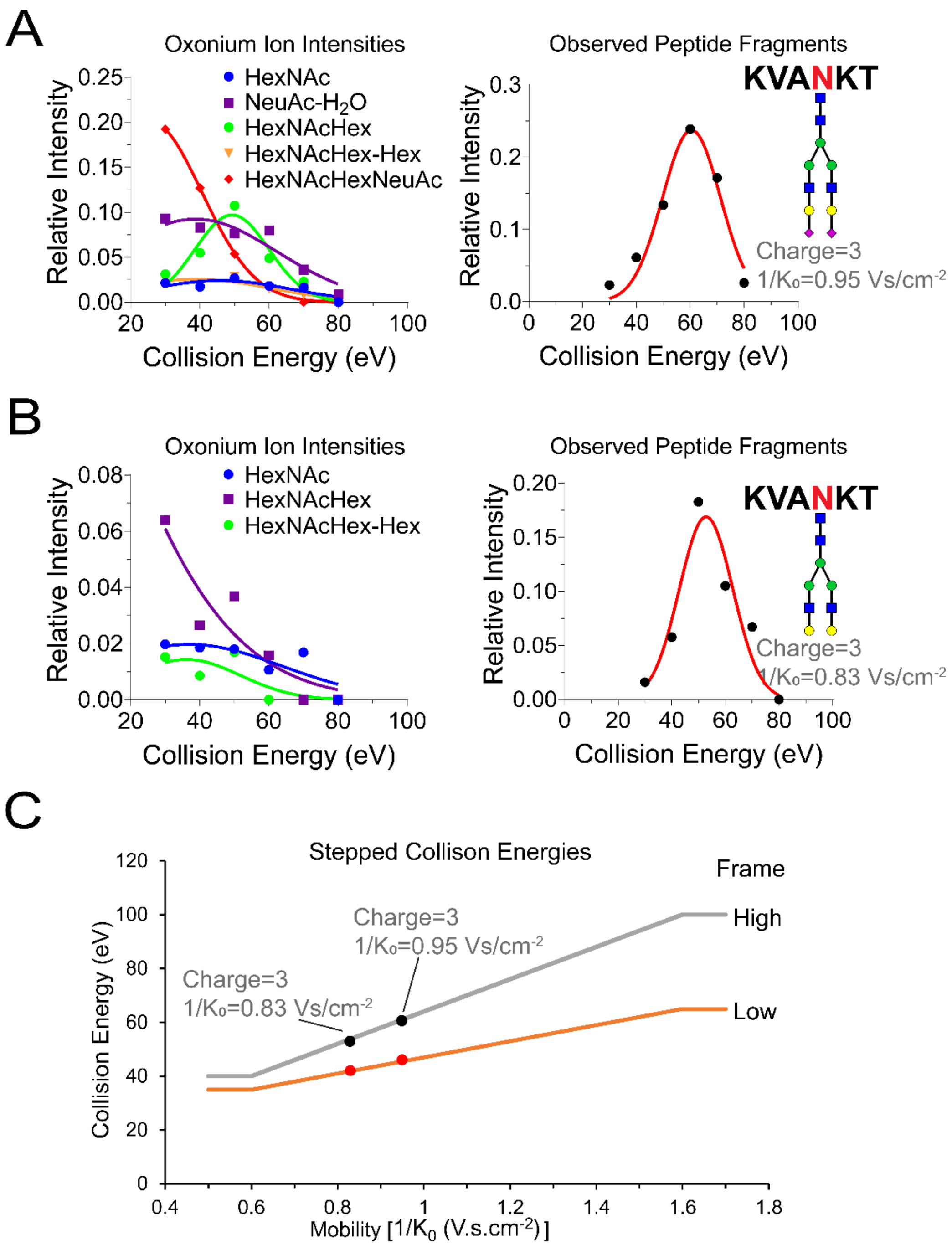


**Figure S6.** Optimized stepped collision energies for optimal glycopeptide detection on timsTOF Pro. Relative intensities of the oxonium ions and peptide fragments (b and y ions) at various collision energies for **(A)** SGP at reduced ion mobility ($1/K_0$) 0.95 for charge state 3 and **(B)** asialo-SGP at reduced ion mobility ($1/K_0$) 0.83 for charge state 3. The optimal collision energy for SGP peptide fragments is at ~ 60 eV and for asialo-SGP at ~ 50 eV, while the optimal collision energies for oxonium ions were ~ 45 and ~ 40 eV, respectively. **(C)** Final stepped collision energies (SCE) calibration curve linearly extrapolated based on these two glycopeptides from reduced ion mobility ($1/k_0$) 0.6 to 1.6, combining high and low energy frames.

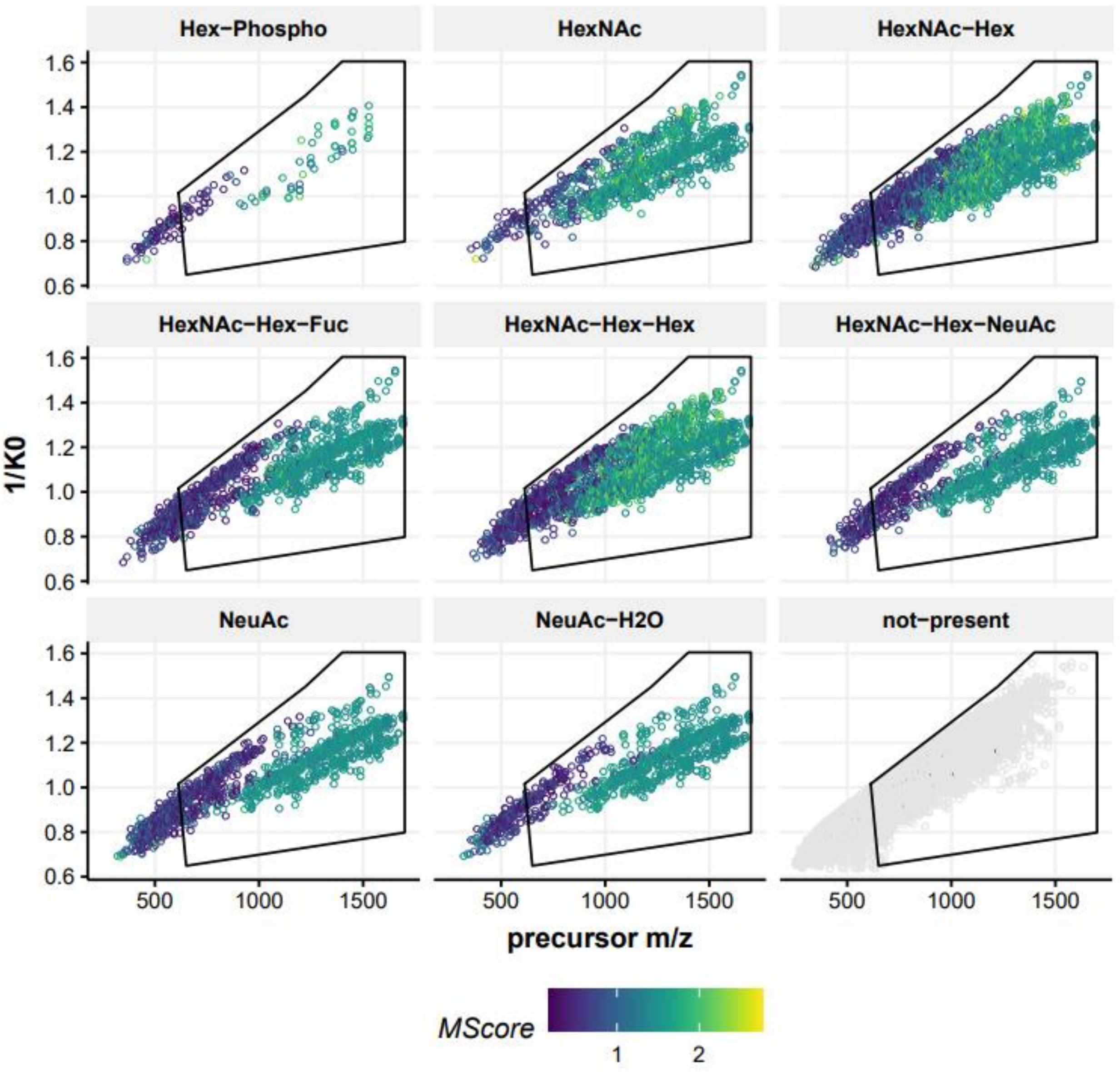


**Figure S7.** Density plot for the distribution for *m/z* vs reduced ion mobility ($1/k_0$) of all the precursor ions containing the glyco-oxonium ions 243. 0270 (Hex-Phospho), 204.0872 (HexNAc), 366.1400 (HexNAc-Hex), 512.198 (HexNAc-Hex-Fuc), 528.198 (HexNAc-Hex-Hex), 657.2354 (HexNAc-Hex-NeuAc), 292.1032 (NeuAc) and 274.0921 (NeuAc-$H_2O$) from human neutrophils. While this black polygon contains all the glycopeptide precursor with M-score > 1.3, precursors with M-score < 1.3 are mostly noise.

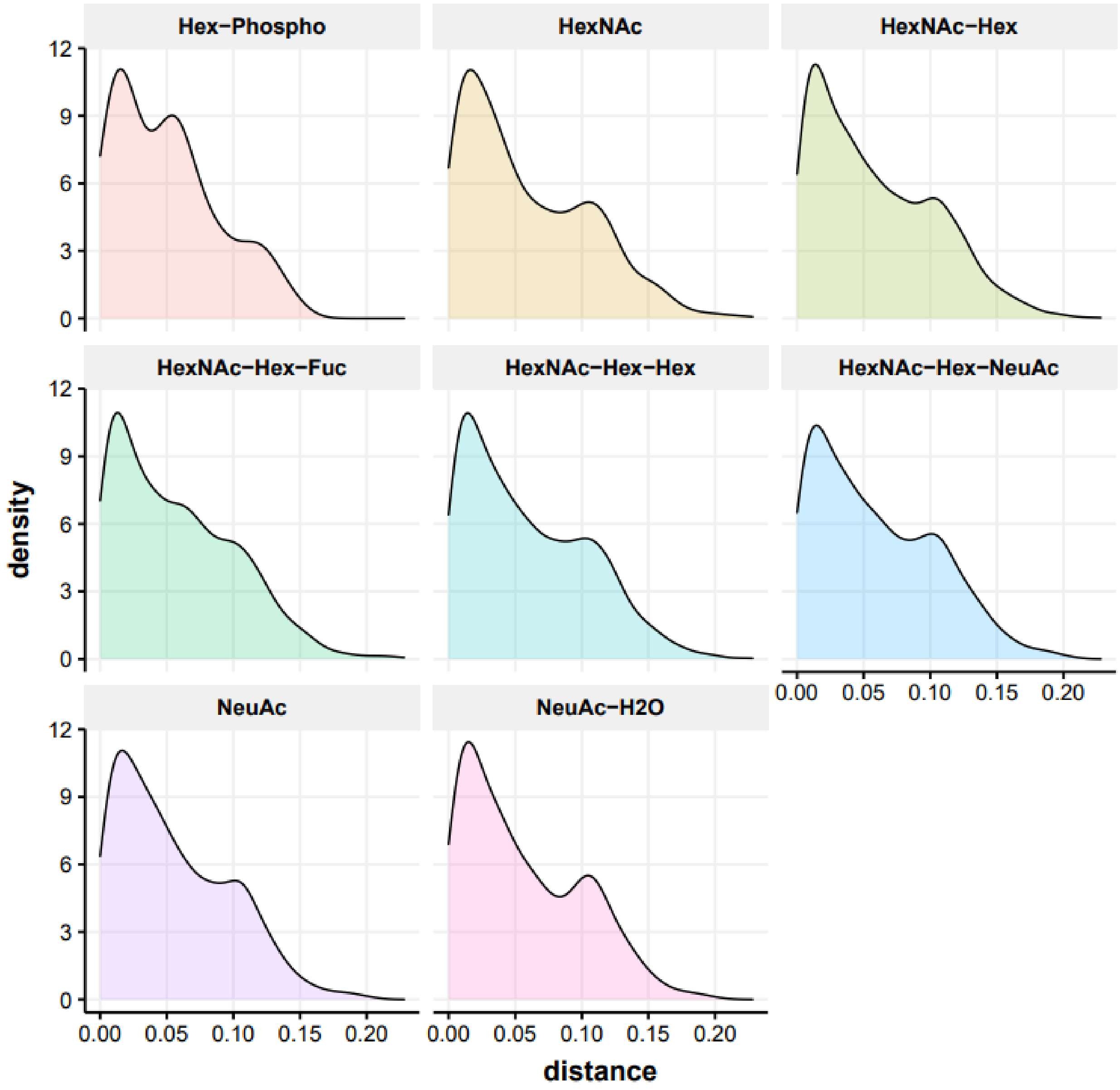


**Figure S8**. Distribution of the glyco-oxonium ion containing glycopeptides from human neutrophils inside the polygon demonstrating different glycopeptides with separate glycan types (high mannose vs sialylated) are indistinguishable by using the timsTOF Pro.

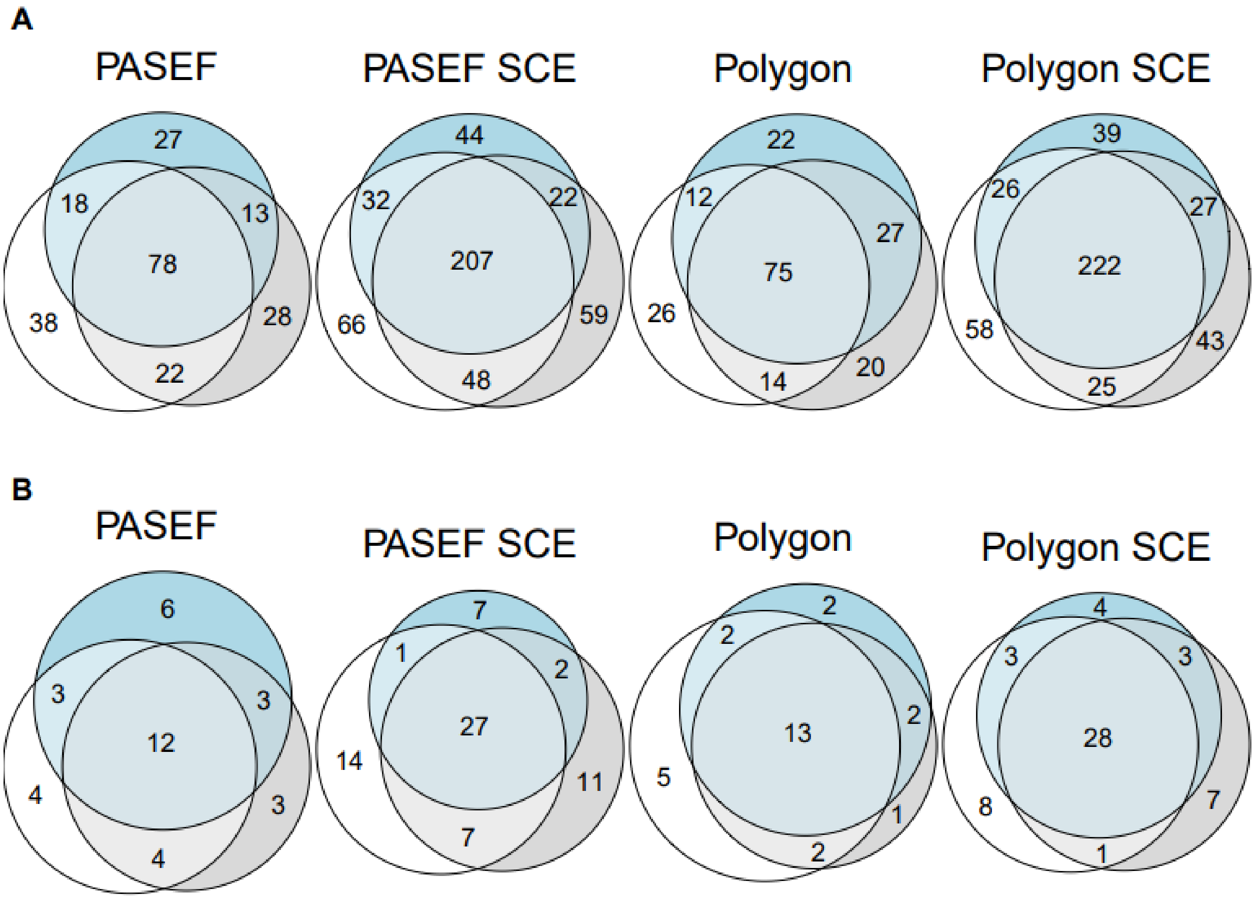


**Figure S9.** Performance comparison of four different methods on the timsTOF Pro for glycoproteomics applied to the neutrophil samples. Overlap of all the annotated **(A)** glycopeptides and **(B)** glycoproteins across three replicated MS/MS measurements in PASEF, PASEF SCE (*i.e* with stepped CE), PASEF with glyco-polygon and PASEF with glyco-polygon and stepped CE. Glyco-polygon demonstrates advantages on glycopeptide identification reproducibility within replicates.

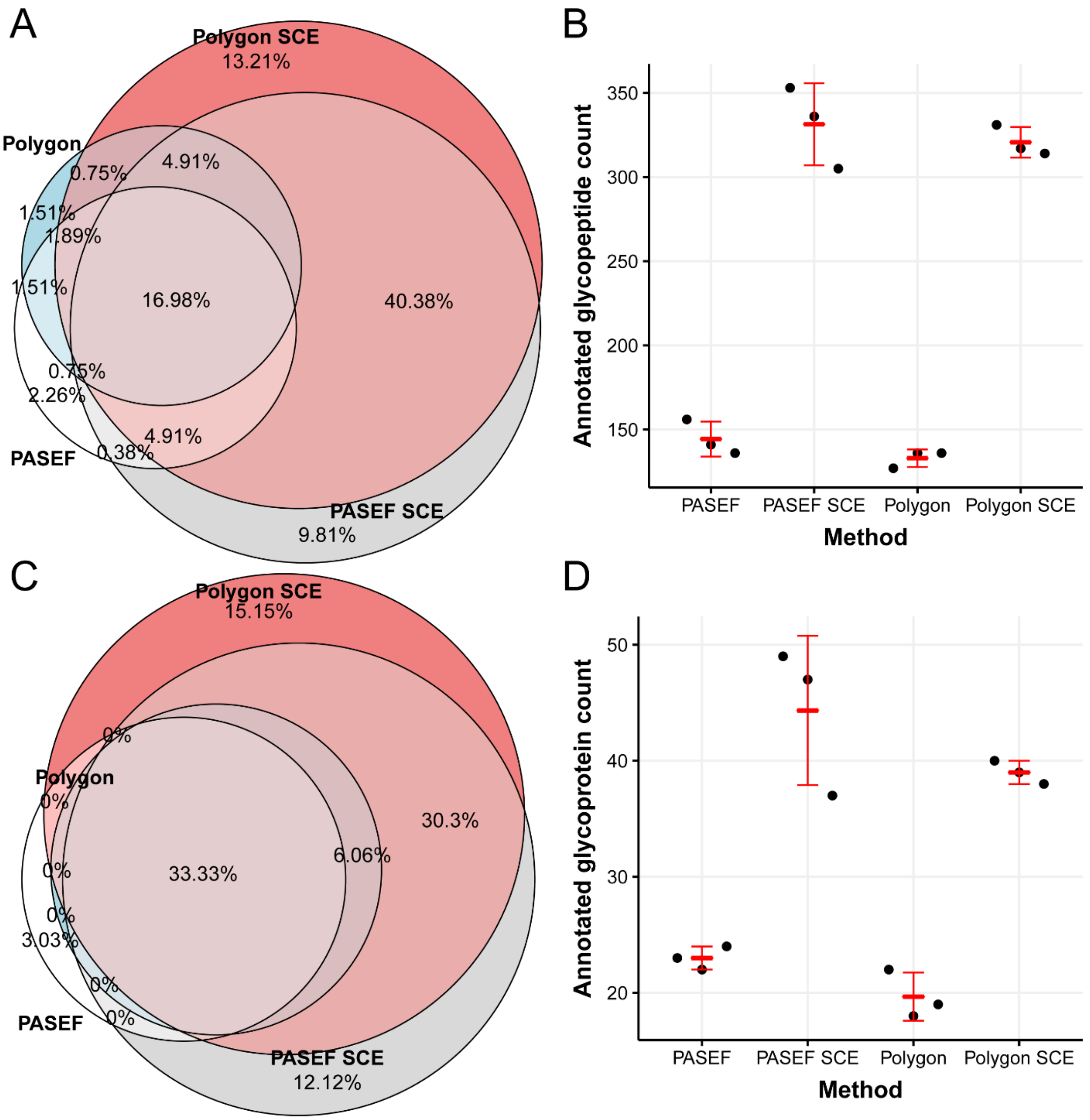


**Figure S10. Performance comparison of four different methods on the timsTOF Pro for glycoproteomics applied to the neutrophil samples**. Overlap of all the **(A)** glycopeptides, **(C)** glycoproteins annotated in all three technical replicates and counts of **(B)** annotated glycopeptides and **(D)** glycoproteins in PASEF, PASEF SCE (*i.e* with stepped CE), PASEF with glyco-polygon and PASEF with glyco-polygon and stepped CE. SCE method demonstrates clear advantages on glycopeptide identification while the IM polygon results in significant improvement of reproducibility across sample replicates.

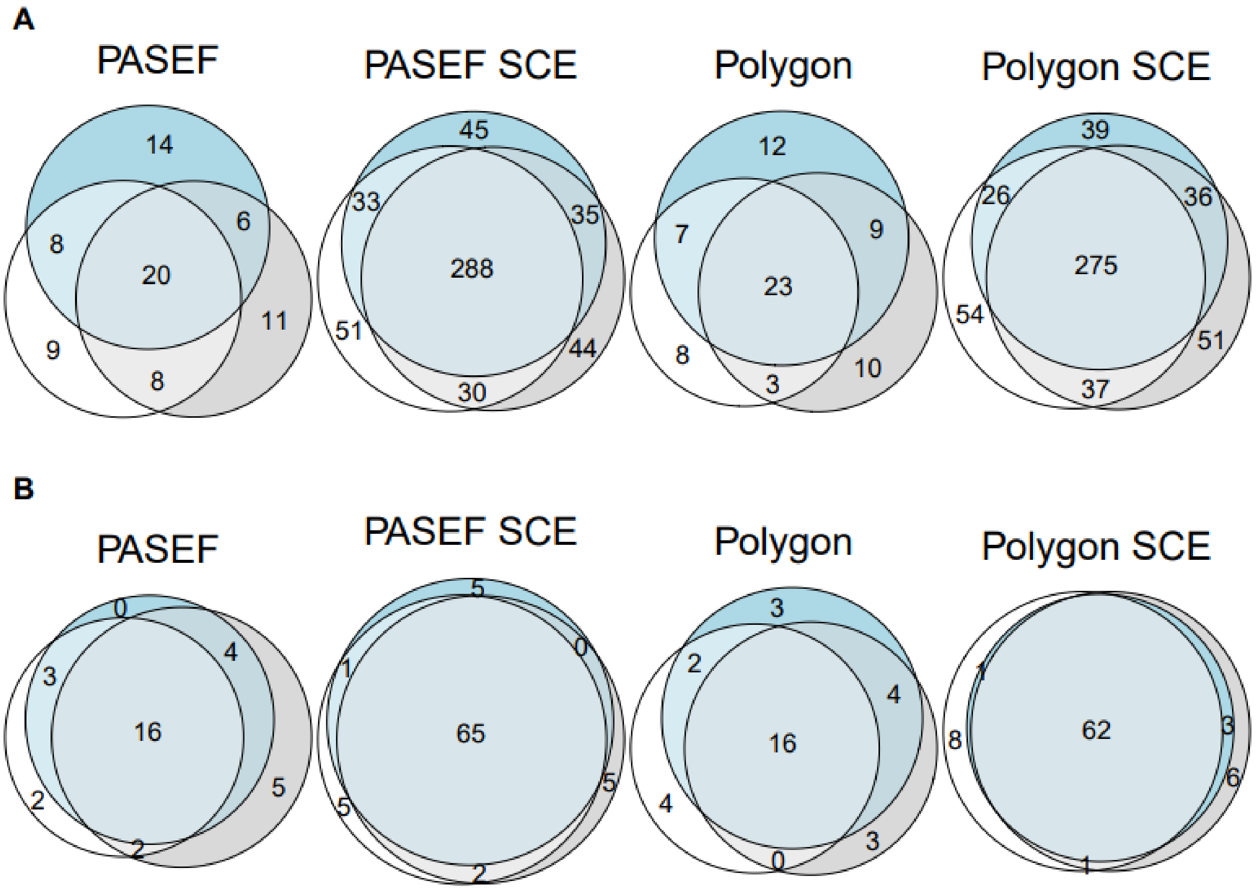


**Figure S11.** Performance comparison of four different methods on the timsTOF Pro for glycoproteomics applied to the plasma samples. Overlap of all the annotated **(A)** glycopeptides and **(B)** glycoproteins across three replicated MS/MS measurements in PASEF, PASEF SCE (*i.e.* with stepped CE), PASEF with glyco-polygon and PASEF with both glyco-polygon and stepped CE. Glyco-polygon demonstrates advantages on glycopeptide identification reproducibility within replicates.

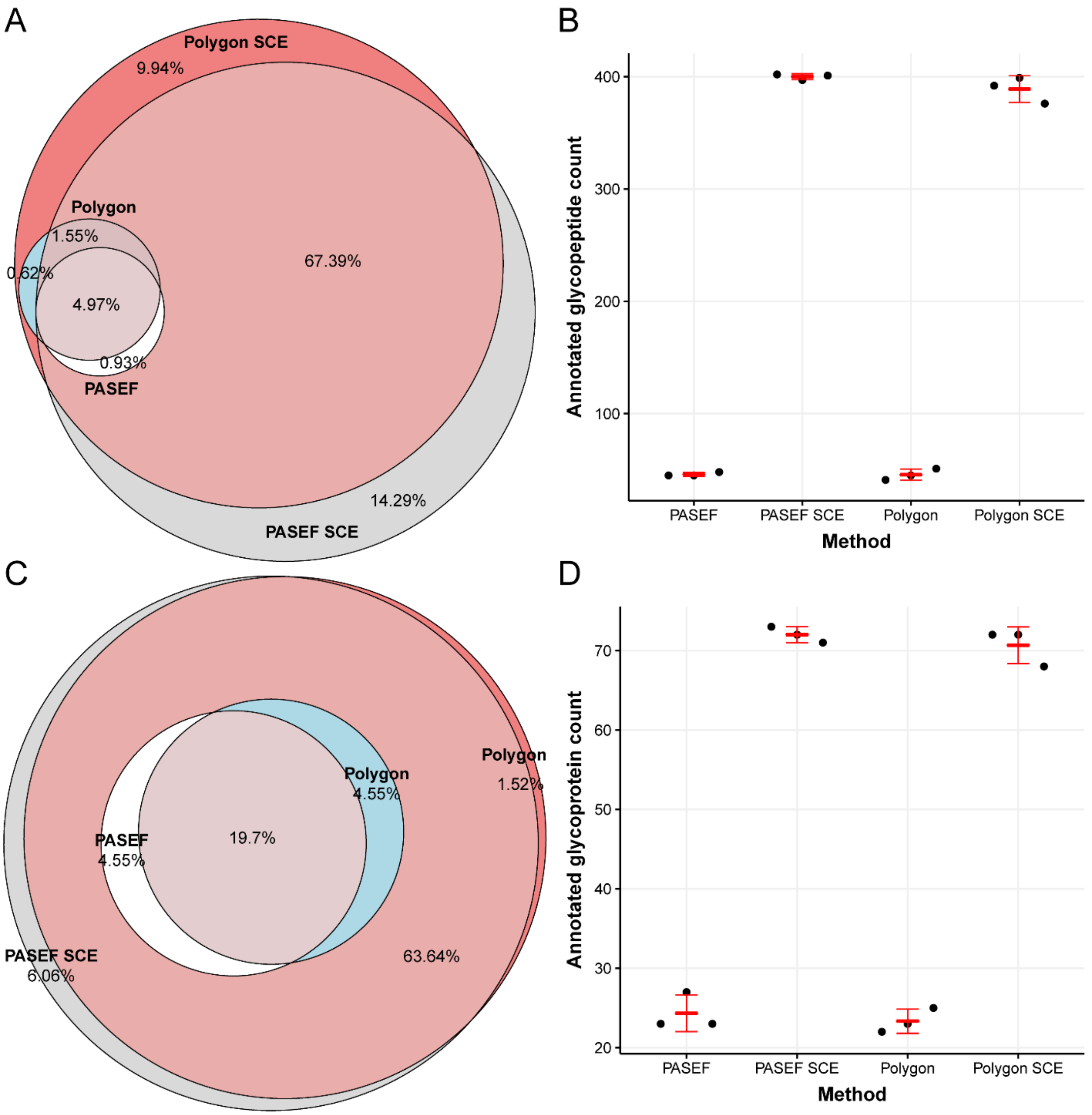


**Figure S12. Performance comparison of four different methods on the timsTOF Pro for glycoproteomics applied to the human plasma sample**. Overlap of all the **(A)** glycopeptides, **(C)** glycoproteins, **(B)** annotated glycopeptides and **(D)** glycoproteins in PASEF, PASEF (ST *i.e* with stepped CE), PASEF with glyco-polygon and PASEF with glyco-polygon and stepped CE. The use of the glyco-polygon demonstrates clear advantages on both glycopeptide identification and glycoprotein coverage.

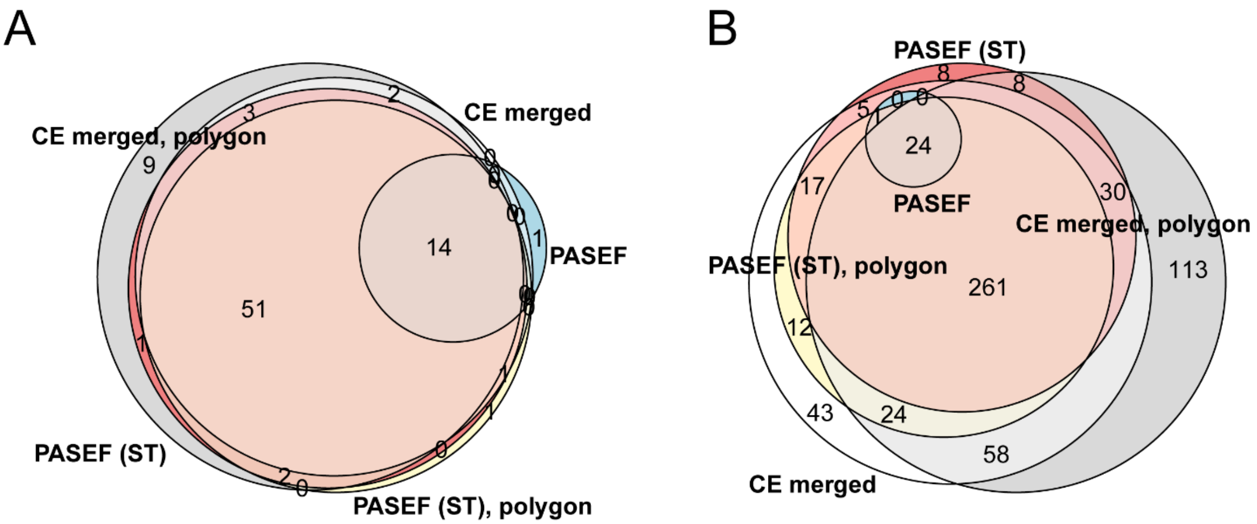


**Figure S13.** Performance comparison of five different methods on the timsTOF Pro for glycoproteomics applied to the human plasma sample. Overlap of all the annotated **(A)** glycoproteins and **(B)** glycopeptides demonstrate clear benefit of using more than two CE for precursor fragmentation during data acquisition.

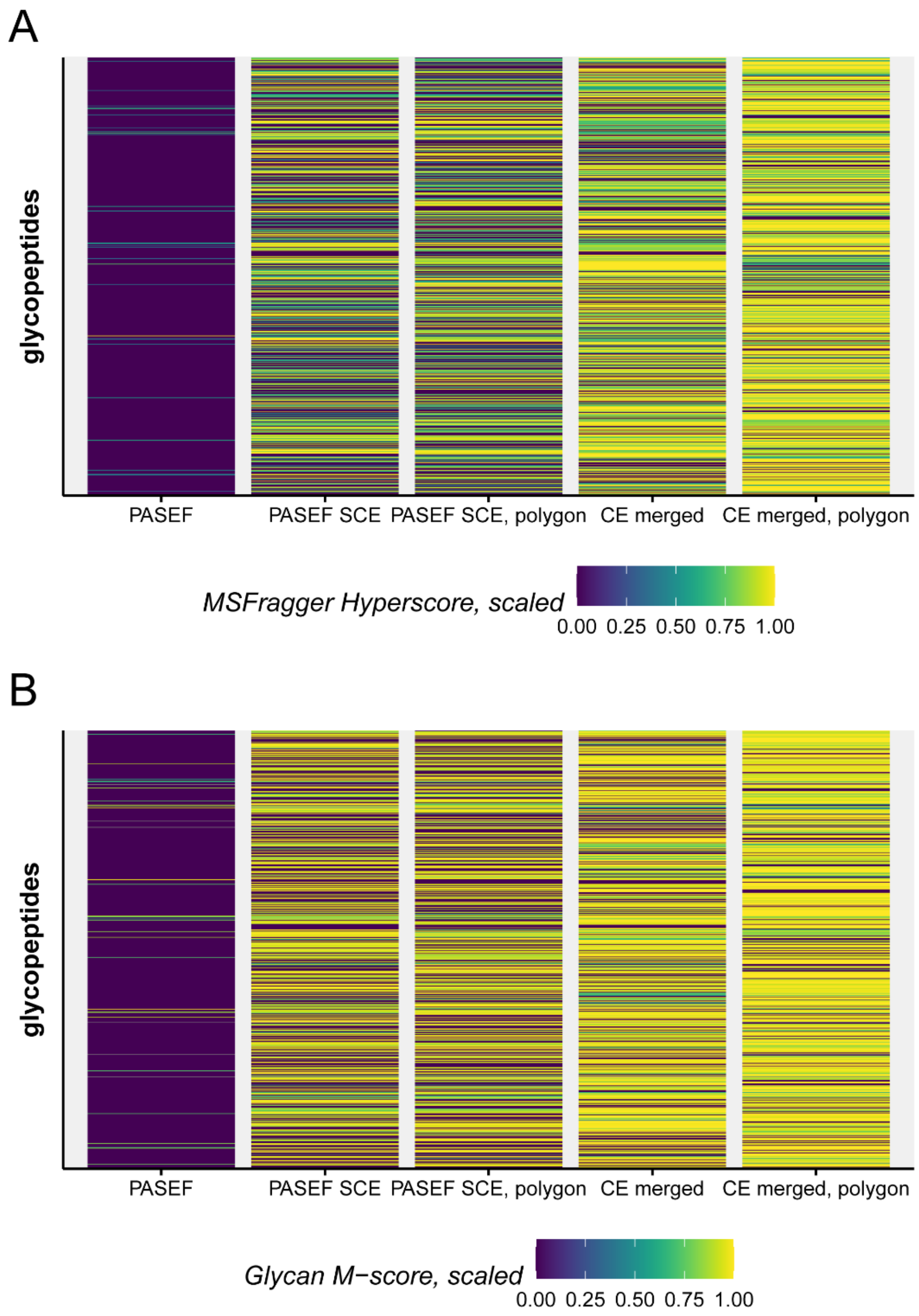


**Figure S14.** Performance comparison of five different methods on the timsTOF Pro for glycoproteomics applied to the human plasma sample. MSfragger peptide annotation score **(A)** and glycan M-score **(B)** both indicate clear increase in data files with CE merged spectra. Scores of each peptide from five different methods are scaled to the maximum value before plotting.

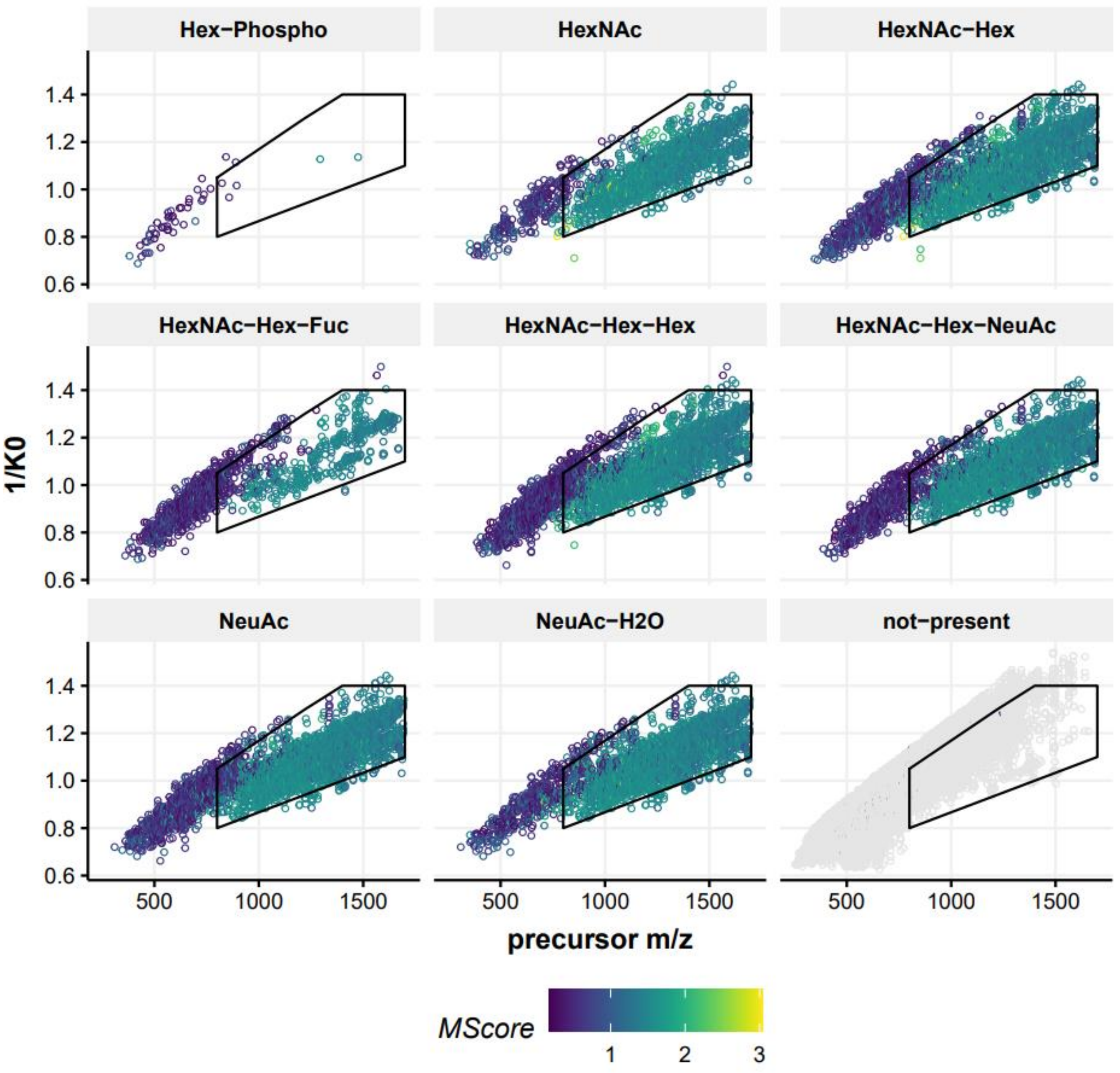


**Figure S15.** Density plot for the distribution for *m/z* vs reduced ion mobility ($1/k_0$) of all the precursor ions containing the glyco-oxonium ions 243.0270 (Hex-Phospho), 204.0872 (HexNAc), 366.1400 (HexNAc-Hex), 512.198 (HexNAc-Hex-Fuc), 528.198 (HexNAc-Hex-Hex), 657.2354 (HexNAc-Hex-NeuAc), 292.1032 (NeuAc) and 274.0921 (NeuAc-$H_2O$) from human plasma. The new stricter black polygon contains all the glycopeptide precursor with M-score > 1.3, describing the ROI for glycopeptide sequencing on the TIMSTOF Pro.

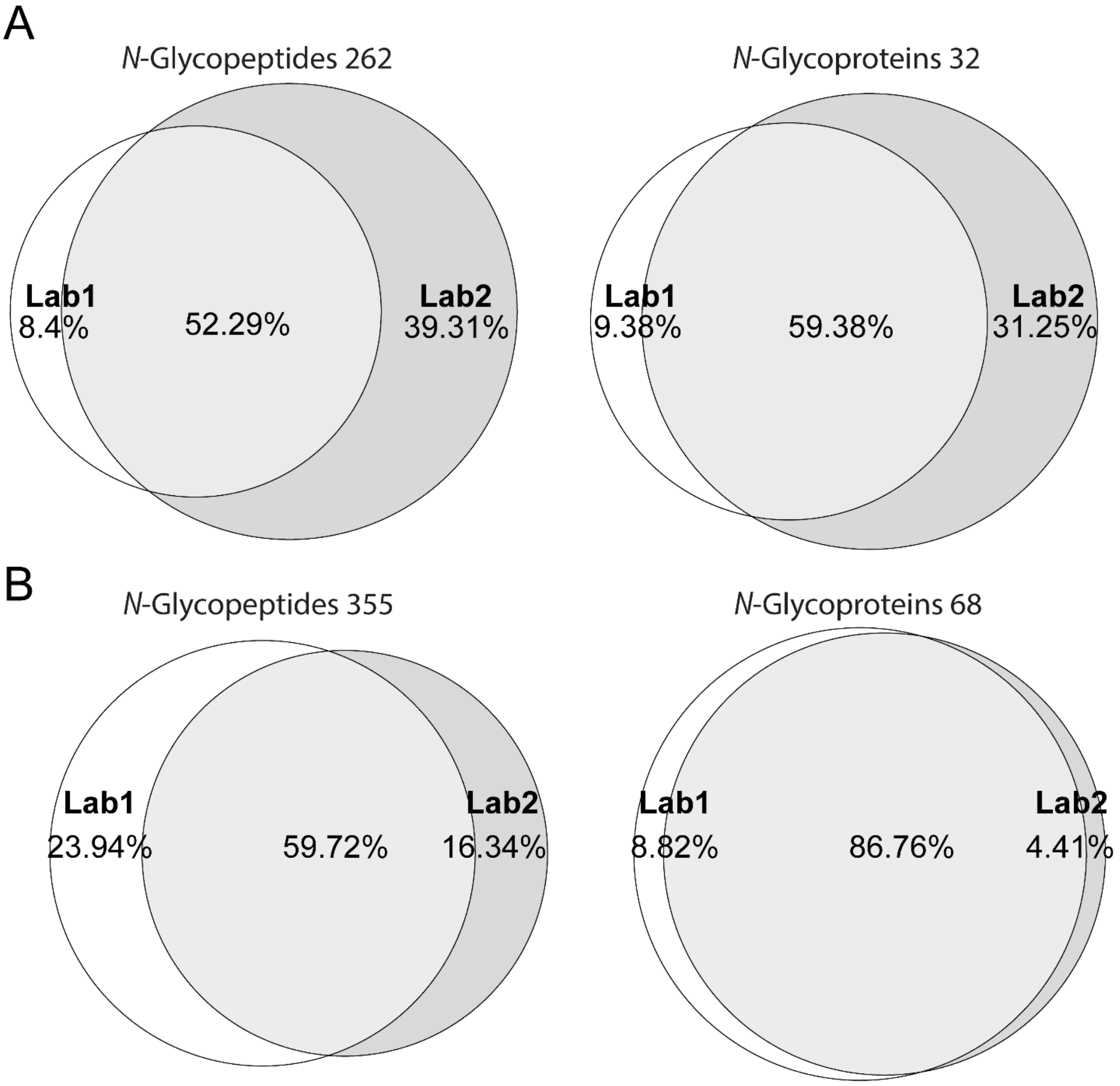


**Figure S16.** Interlaboratory comparison of the oxonium ion-guided ion mobility-assisted glycoproteomics workflow on the timsTOF Pro. Overlap of the *N*-glycopeptides and *N*-glycoproteins in the **(A)** human neutrophil and **(B)** human plasma sample that was shared between two different laboratories and analyzed using the SCE PASEF glyco-polygon method in triplicates. Only *N*-glycopeptides and *N*-glycoprotein present in all three replicates were used to compare the interlaboratory reproducibility.

**Table S1. Performance comparison of different methods of the timsTOF Pro for glycoproteomics applied to the human neutrophil sample.**

| Method | Replicate | Glyco psms | Unique glycopeptides | Unique glycoproteins |
|---|---|---|---|---|
| PASEF | 1 | 247 | 156 | 23 |
| PASEF | 2 | 219 | 141 | 22 |
| PASEF | 3 | 205 | 136 | 24 |
| PASEF polygon | 1 | 216 | 127 | 22 |
| PASEF polygon | 2 | 225 | 136 | 18 |
| PASEF polygon | 3 | 206 | 136 | 19 |
| PASEF SCE | 1 | 609 | 353 | 49 |
| PASEF SCE | 2 | 570 | 336 | 47 |
| PASEF SCE | 3 | 512 | 305 | 37 |
| PASEF SCE polygon | 1 | 590 | 331 | 40 |
| PASEF SCE polygon | 2 | 541 | 317 | 39 |
| PASEF SCE polygon | 3 | 531 | 314 | 38 |

**Table S2. Performance comparison of different methods of the timsTOF Pro for glycoproteomics applied to the human plasma sample.**

| Method | Replicate | Glyco psms | Unique glycopeptides | Unique glycoproteins |
|---|---|---|---|---|
| PASEF | 1 | 77 | 45 | 23 |
| PASEF | 2 | 76 | 45 | 27 |
| PASEF | 3 | 80 | 48 | 23 |
| PASEF polygon | 1 | 74 | 41 | 22 |
| PASEF polygon | 2 | 79 | 45 | 23 |
| PASEF polygon | 3 | 97 | 51 | 25 |
| PASEF SCE | 1 | 895 | 402 | 73 |
| PASEF SCE | 2 | 868 | 397 | 72 |
| PASEF SCE | 3 | 872 | 401 | 71 |
| PASEF SCE polygon | 1 | 889 | 392 | 72 |
| PASEF SCE polygon | 2 | 874 | 399 | 72 |
| PASEF SCE polygon | 3 | 844 | 376 | 68 |

**Table S3.Performance comparison of different methods and CE spectra summing on the timsTOF Pro for glycoproteomics applied to the human plasma sample**.

| Method | Glyco psms | Unique glycopeptides | Unique glycoproteins |
|---|---|---|---|
| PASEF | 51 | 29 | 15 |
| PASEF SCE | 788 | 368 | 72 |
| PASEF SCE, polygon | 820 | 378 | 69 |
| CE merged | 1052 | 478 | 71 |
| CE polygon merged | 1259 | 545 | 82 |

**Table S4. Performance comparison of PASEF SCE method with/without strict glyco-polygon of the human plasma sample with different chromatography gradients.**

| Method, LC gradient length, min | Replicate | Glyco psms | Unique glycopeptides | Unique glycoproteins |
|---|---|---|---|---|
| Without polygon, 15 | 1 | 97 | 77 | 32 |
| Without polygon, 15 | 2 | 92 | 69 | 28 |
| Without polygon, 15 | 3 | 99 | 73 | 25 |
| Without polygon, 30 | 1 | 276 | 185 | 51 |
| Without polygon, 30 | 2 | 285 | 189 | 48 |
| Without polygon, 30 | 3 | 275 | 183 | 47 |
| Without polygon, 60 | 1 | 574 | 332 | 65 |
| Without polygon, 60 | 2 | 560 | 328 | 63 |
| Without polygon, 60 | 3 | 582 | 348 | 67 |
| Without polygon, 90 | 1 | 667 | 376 | 67 |
| Without polygon, 90 | 2 | 678 | 367 | 67 |
| Without polygon, 90 | 3 | 700 | 384 | 69 |
| Polygon, 15 | 1 | 162 | 112 | 37 |
| Polygon, 15 | 2 | 160 | 108 | 36 |
| Polygon, 15 | 3 | 155 | 110 | 36 |
| Polygon, 30 | 1 | 438 | 271 | 63 |
| Polygon, 30 | 2 | 441 | 276 | 57 |
| Polygon, 30 | 3 | 470 | 273 | 55 |
| Polygon, 60 | 1 | 732 | 381 | 69 |
| Polygon, 60 | 2 | 758 | 393 | 74 |
| Polygon, 60 | 3 | 729 | 388 | 68 |
| Polygon, 90 | 1 | 873 | 454 | 74 |
| Polygon, 90 | 2 | 893 | 451 | 74 |
| Polygon, 90 | 3 | 905 | 451 | 74 |
